# Spatially specific, closed-loop infrared thalamocortical deep brain stimulation

**DOI:** 10.1101/2023.10.04.560859

**Authors:** Brandon S Coventry, Georgia L Lawlor, Christina B Bagnati, Claudia Krogmeier, Edward L Bartlett

**Affiliations:** Weldon School of Biomedical Engineering, Purdue University, West Lafayette, IN USA; Center for Implantable Devices and the Institute for Integrative Neuroscience, Purdue University, West Lafayette, IN USA; Department of Computer Graphics Technology, Purdue University, West Lafayette, IN USA; Department of Biological Sciences, Purdue University, West Lafayette, IN USA

**Keywords:** Infrared Neural Stimulation, Deep Brain Stimulation, Closed-Loop DBS, Deep Reinforcement Learning

## Abstract

Deep brain stimulation (DBS) is a powerful tool for the treatment of circuitopathy-related neurological and psychiatric diseases and disorders such as Parkinson’s disease and obsessive-compulsive disorder, as well as a critical research tool for perturbing neural circuits and exploring neuroprostheses. Electrically-mediated DBS, however, is limited by the spread of stimulus currents into tissue unrelated to disease course and treatment, potentially causing undesirable patient side effects. In this work, we utilize infrared neural stimulation (INS), an optical neuromodulation technique that uses near to mid-infrared light to drive graded excitatory and inhibitory responses in nerves and neurons, to facilitate an optical and spatially constrained DBS paradigm. INS has been shown to provide spatially constrained responses in cortical neurons and, unlike other optical techniques, does not require genetic modification of the neural target. We show that INS produces graded, biophysically relevant single-unit responses with robust information transfer in thalamocortical circuits. Importantly, we show that cortical spread of activation from thalamic INS produces more spatially constrained response profiles than conventional electrical stimulation. Owing to observed spatial precision of INS, we used deep reinforcement learning for closed-loop control of thalamocortical circuits, creating real-time representations of stimulus-response dynamics while driving cortical neurons to precise firing patterns. Our data suggest that INS can serve as a targeted and dynamic stimulation paradigm for both open and closed-loop DBS.

**Significance Statement:** Despite initial clinical successes, electrical deep brain stimulation (DBS) is fraught with off-target current spillover into tissue outside of therapeutic targets, giving rise to patient side effects and the reduction of therapeutic efficacy. In this study, we validate infrared neural stimulation (INS) as a spatially constrained optical DBS paradigm by quantifying dose-response profiles and robust information transfer through INS driven thalamocortical circuits. We show that INS elicits biophysically relevant responses which are spatially constrained compared to conventional electrical stimulation, potentially reducing off-target side effects. Leveraging the spatial specificity of thalamocortical INS, we used deep reinforcement learning to close the loop on thalamocortical INS and showed the ability to drive subject-specific thalamocortical circuits to target response states in real time.

## Introduction

Electrical stimulation of the nervous system has emerged as a potent tool for the treatment and study of a wide variety of neurological diseases(1–5), as well as a key research tool for modulating and mapping neural circuits(6–9). The most prominent of these stimulation paradigms are cochlear implants (CI), which induce sound percepts in individuals with profound hearing loss, and deep brain stimulation (DBS), which has proven effective in treating movement-related symptoms associated with Parkinson’s disease and essential tremor. Additionally, diseases treated by electrical neuromodulation are expanding, with DBS having recently received an FDA humanitarian device exemption for the treatment of obsessive-compulsive disorder while also in clinical trial for major depressive disorder(3), Tourette’s syndrome(10), and epilepsy(1). Peripheral nerve electrical stimulation technologies are also maturing into viable clinical tools, including vagus nerve stimulation for the treatment of epilepsy(11) and carotid sinus stimulation for the treatment of heart disease(2).

Despite initial clinical success, electrical paradigms of neuromodulation are fraught with undesirable current spillover into off-target neural circuits(12–16) leading to undesirable side effects and a reduction in therapeutic efficacy(17–19). The development of focal stimulation strategies is paramount to more effective clinical stimulation and the improvement of patient side-effect profiles. One such tool is infrared neural stimulation (INS), an optical modality which stimulates nerves and neurons using near to mid infrared wavelength (700-2000 nm) light(20–23). INS has shown spatially specific recruitment of both peripheral nerves(16, 24, 25) and central neurons(22, 23). Importantly, INS does not require genetic manipulation necessary for other optical stimulation methods(26), acting purportedly on intrinsic cell biophysics(27). INS also shows promising safety profiles for translation to human patients(28–30) and has found use in diagnostic targeting of human nerve roots in surgical resection procedures(31). While INS is a promising modality for neuromodulation therapies, progress towards optically-based DBS (oDBS) is hindered by a lack of understanding of INS entrainment of thalamocortical and sub-thalamocortical networks; the understanding of which is necessary for treating “circuitopathies” associated with diseases treated by DBS(32–37). Specifically, there is a dearth of information related to dose-response dependencies of INS laser parameters in circuital recruitment and the resulting spread of activation across neural circuits.

In this study, we validate INS as a potent oDBS paradigm by quantifying INS dose-response profiles from varying laser parameters, INS driven information transmission across the thalamocortical synapse, and spatial specificity of network INS in the rat auditory thalamocortical model. Our experiments show strong evoked firing rate dependence on applied laser energy with increases in thalamocortical information transfer with increased laser energy. We further show that INS evokes cortical activity that maintains typical thalamocortical response profiles with constrained spread of activation well below the spread of electrical stimulation. Owing to the targeted neural activation of INS, we engineered a closed-loop control approach called SpikerNet, a deep reinforcement learning (RL) based reactive DBS system(38, 39). Closed-loop DBS utilizes feedback from biomarkers of disease to apply stimulation only when needed(40) and has shown advantageous in therapeutic efficacy and battery life(41). However, the relatively simple control algorithms of conventional closed-loop DBS limit the ability to capture complex dynamics of neural activity related to disease which can cause interference with normal activity, such as interruption of volitional movement(42) which is further exacerbated by large scale activation from electrical stimulation(43). More complex control methods are advantageous in accounting for brain wide state changes, such as sleep wake cycles(44). We therefore utilized deep RLs ability to develop statistical mappings of systems in response to state perturbations in order to drive cortical activity to desired firing states.

## Results

### Implanting Stimulation and Recording Devices

The auditory pathway has a rich history of neuromodulation, with electrical stimulation of the cochlea resulting in cochlear implants, one of the first and most successful clinical neuromodulation devices(45). Other clinical auditory devices include the auditory brainstem and midbrain implants(46, 47) with electrical neuromodulation across all auditory nuclei(48–50) are being investigated for clinical viability. Auditory thalamocortical circuits are particularly suited for study because the regional architecture of the auditory thalamus permits stimulation of both core and belt pathways in rodents, primates(51), and humans(52) using a single dorsoventrally oriented electrode. This enables testing stimulation strategies simultaneously in both tonotopic core pathways and higher-order belt pathways, along with the ability to rapidly test circuit function with minimally invasive scalp evoked auditory potentials(53–55) before and after device implantation. To facilitate understanding of dose-response effects of network function elicited through INS, rats were implanted with fiber optic optrodes into the medial geniculate body of the auditory thalamus (MGB). The ventral and dorsal divisions of the MGB have primary excitatory afferents to layer 3/4 of auditory cortex(56) (Fig 1A,C). Sixteen channel planar arrays were implanted into layer 3/4 of primary auditory cortex (Fig 1A), and all MGB subdivisions have at least some projection to primary auditory cortex(57). Postmortem histological analyses confirmed placement of optrodes into the MGB (Fig 1D, Supplementary Methods).

**Figure 1.**
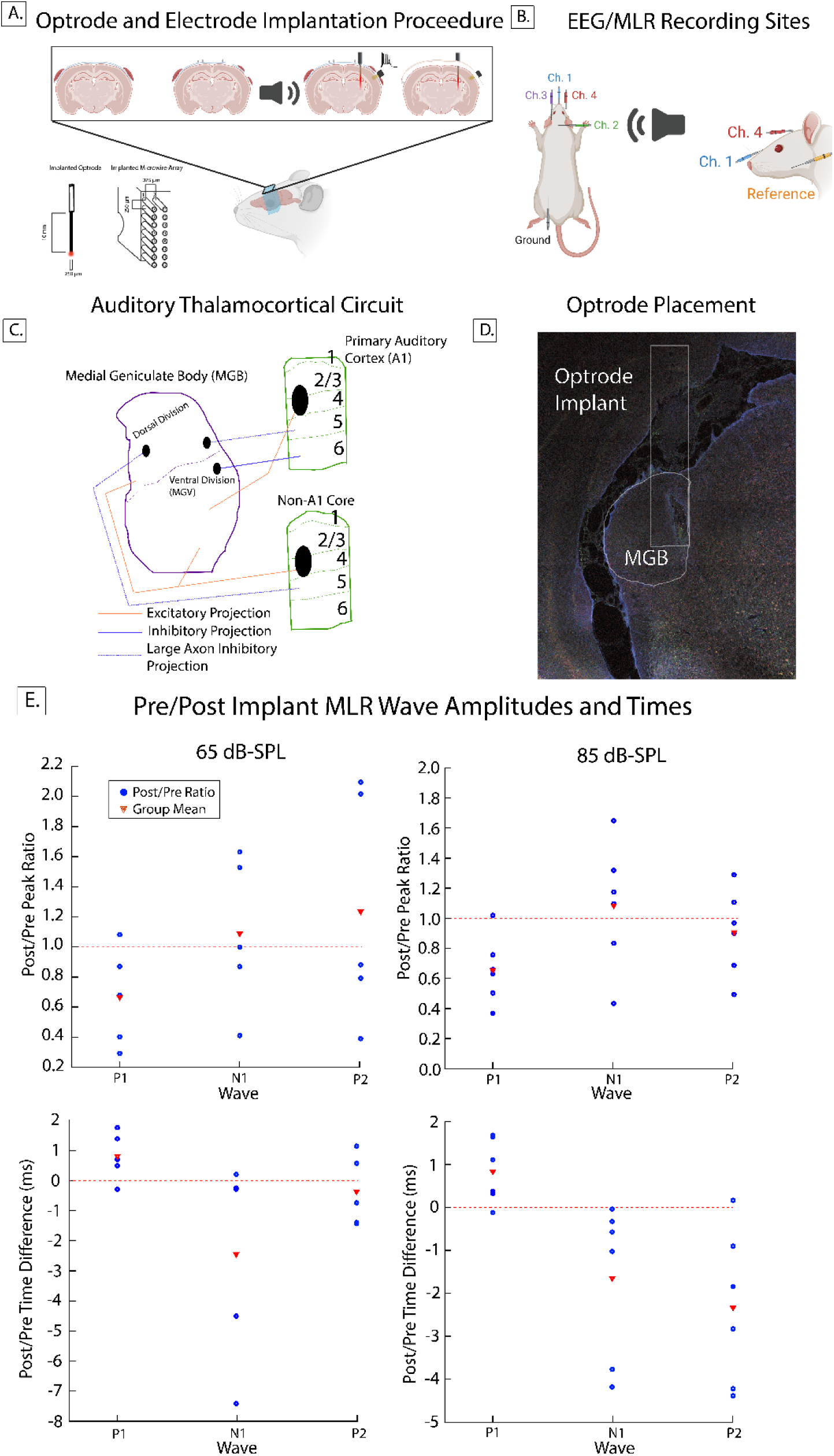
Implantation and EEG-MLR procedures. A. Left: Rodents were implanted with fiber optic optrodes into the medial geniculate body and 16 channel microwire arrays into auditory cortex. Placement of microwire array was confirmed by tonic single unit responses evoked from 80 dB filtered Gaussian noise stimuli during implantation. A. Right: Schematic of the 4-channel EEG-MLR recording preparation. B. Left: Schematic of the rodent auditory thalamocortical circuit. Stimulation optrodes were placed in the ventral division of the medial geniculate body with primary excitatory efferent projections to layer 3-4 of primary auditory cortex. Microwire array recording electrodes were placed in layer 3-4 of primary auditory cortex confirmed during surgery by low-latency single unit activity. B. Right: Histological images demonstrate placement of stimulation optrode was within medial geniculate body. C. EEG-MLR pre-post surgical ratios show small changes in wave P1, N1, and P2 correlates of auditory thalamocortical function in amplitude and latency due to passive presence of device at 65 or 85 dB-SPL click stimuli. While changes in amplitudes and latencies were observed effects, differences did not rise to level of significance (p>0.05). Rodent implantation and EEG diagrams were created using BioRender under publication license.

Development of INS into a clinically viable neuromodulation system has been limited by a lack of understanding of underlying stimulation mechanisms and stimulus to response mappings. A confounding factor is that commercial INS systems are not widely available and are prohibitively expensive or removed from the market by product recalls (58). To facilitate continued INS studies, we developed INSight, a low-cost open source INS and optical stimulation system which uses off the shelf components for ease of building and modification. Importantly, INSight can integrate into established recording systems. Materials, build instructions, and calibrations are found in the supplementary material (Fig S10-11) and the INSight Github repository: https://github.com/bscoventry/INSight.

### Changes in neural activity due to presence of devices in the brain

Implantation of recording and stimulation devices present a critical assault to normal neural function(59, 60). Therefore, we first considered the effect of the presence of stimulation and recording devices in brain activity through auditory evoked mid-latency responses (MLRs) in a subset of rats (n=6). MLRs stimuli consisted of evoked responses to auditory click trains with recordings taking place 24 hours before and 72 hours after implantation procedures. MLRs report auditory generators in thalamus and cortex and serve as a read out of neural ensemble function(61–64). We utilized a 4 positive channel EEG recording configuration to allow for responses of thalamocortical generators and from rostral brainstem regions(53)(Fig 1A Right) on each hemisphere and we analyzed ratios of post-pre positive peaks 1 and 2 (P1,P2) corresponding to brainstem and cortical generators, respectively and negative peak 1 (N1 or N1-P2) corresponding to thalamic generators (Fig 1B). While there was some variability in wave amplitudes and latencies, comparisons of evoked activity resulting from click-train auditory stimuli at 65 and 85 dB-SPL (Fig 1E) showed no significant difference in response (p>0.05, Wilcoxon sign-rank) suggesting that presence of stimulation optrodes and recording electrodes did not significantly damage or alter thalamic and cortical activity at the onset of INS experiments. It should be noted that post-surgical recordings were performed 72 hours after surgery, well within the device heal-in window(65) with further neural reorganization likely to occur throughout the duration of the study.

### Dose-response relationships of cortical neuron response from thalamic INS

We next examined the interplay of INS laser energy and interstimulus pulse intervals (ISI) on evoked cortical single unit firing rates. Excitatory peristimulus time histograms (PSTHs) of single units which were responsive to INS stimuli (Z-score increase ≥ 7.84 from basal firing rate, *p* < 0.00001) were analyzed. Units showing inhibitory responses or no change from basal firing rates were excluded from the present study. While INS dose-response relationships have been studied in cortex(22, 23), they remain unstudied across thalamocortical networks. Dose-response profiles were modeled as a Bayesian linear random effects regression model, allowing us to account for hierarchical structure of data consisting of variability within and between subjects across implantation lifetimes. Bayesian inference is particularly powerful for this model as it provides complete quantification of posterior distributions over all regression parameters and allows for direct uncertainty quantification of parameters. Inference was performed directly on observed data posterior distributions. As Bayesian methods require specification of prior probability distributions for inference, broad, non-informative normal prior distributions were used in inference models. Prior sensitivity analyses were performed in order to ensure prior distributions did not unduly influence inference (Fig S5, Table S2). Dose-response regression models took the form of

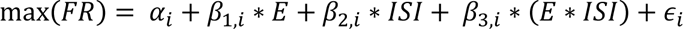

with response variable FR representing the natural log transformed evoked firing rate and independent variables E and ISI being the natural log transformed energy per pulse and inter-stimulus interval respectively. Natural log transformations of response and independent variables were chosen as model comparisons and sensitivity analyses dictated that these models best fit observed data (Fig S5). An error term of *ϵ* was added for uncertainty quantification. Full model descriptions and sensitivity analyses are provided in SI:Bayesian model description (Fig S1-S9). Regression parameters were summarized by their maximum *a priori* estimate (ie most probable value) with independent variables considered significant contributors to response if the highest density interval (HDI) of the parameter distribution corresponding to the 95% most probable parameter values did not overlap 0, following Bayesian inference convention(66). Regression models (Fig 2B) show that INS responsive units had a basal firing rate greater than 0 (*α* MAP = 2.2, 95% HDI exludes 0) with max evoked firing rates depending significantly on applied laser energy (*β*_1_, MAP = 0.58, 95% HDI excludes 0) but not on ISI (*β*_2_, MAP = -0.055) or energy-ISI interactions (*β*_3_, MAP = 0.028). However, the relatively wide spread of the ISI parameter *β*_2_ across 0 suggests a potential critical point in ISI timing past which thalamocortical neurons are unable to entrain to individual pulses and instead integrate INS pulses into a single network event.

**Figure 2.**
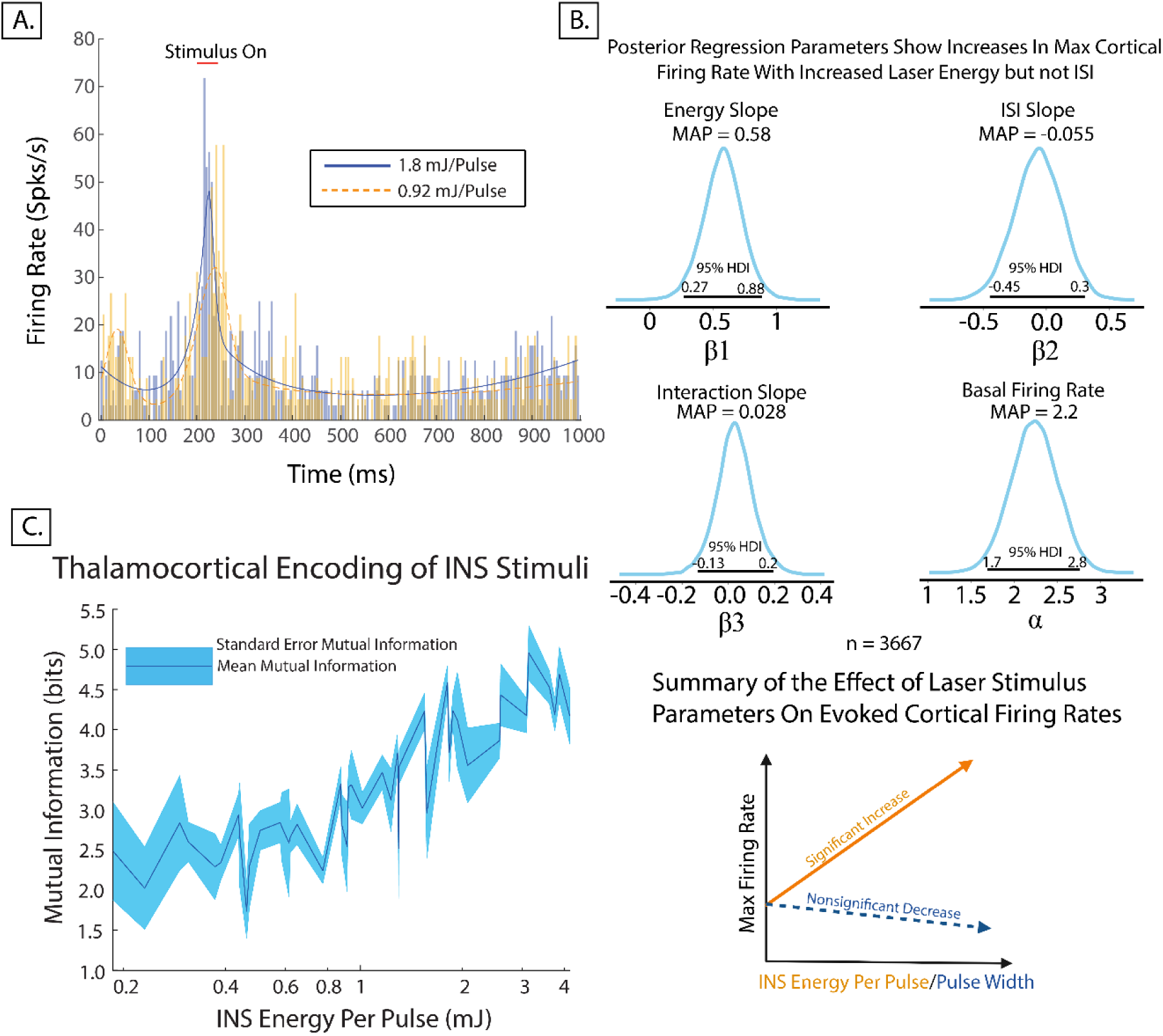
A. Example INS-evoked peristimulus time histograms. BARS estimates of 1.8mJ (solid blue line) and 0.92mJ (dotted orange line) show higher energy pulses drive higher firing, lower latency responses. B. Bayesian hierarchical linear regression models of cortical dose-response profiles elicited from varying INS parameters. Distributions of regression parameters are given for applied laser energy, laser pulse width, and laser energy-pulse width interactions. Regressions show that increases in applied energy significantly increase maximum cortical firing rates with a maximum a priori estimate 0.58 increase in log firing rate in response to increases in log energy (95% HDI does not overlap 0). The width of the 95% HDI of the energy parameter (0.27-0.88) suggests that while cortical firing rates increase with increases in laser energy, INS dose-response profiles are dependent on the physiology of the neuron. Slight decreases in firing rate with increased laser pulse widths were observed (MAP = -0.055), but not significant (95% HDI overlaps 0). Laser energy and pulsewidth interactions also did not significantly change evoked cortical firing rates (95% HDI overlaps 0). Basal firing rates of neurons were significantly above zero (95% HDI does not overlap 0, MAP estimate = 2.2). C. Evoked single unit spike train information increases as INS energy increases.

### Cortical encoding of INS stimuli

We next used Shannon mutual information measures [*I*(*R*; *S*_*x*_), *Eq*. 2] to assess and quantify information carried by evoked spike-trains in response to INS stimulation energy. Mutual information measures the reduction of uncertainty in neural response given knowledge of the stimulus. Higher values of information represent more unique and separable encoding of neural response distributions for each stimulus. Stimulus-information profiles were calculated from 5 ms binned estimates of response probability mass distributions during INS conditioned on applied energy. Bias in mutual information resulting from incomplete knowledge of population response distributions was estimated and corrected using the methods of quadratic extrapolation(67, 68). We found that increasing INS energy per pulse resulted in increases in information contained in response spike trains (Fig 2.C). Increases in information are also positively correlated with increased INS energy per pulse showing strong dependence of evoked PSTHs on laser energy, particularly > 0.8 mJ/pulse (Fig. 2C).

Auditory thalamocortical circuits perform complex transformations of inputs at the auditory thalamocortical synapse(69) with cortical neurons employing differential coding strategies across local heterogeneous cells and circuits(70, 71). Therefore, it is imperative that any stimulation modality be able to drive naturalistic response profiles. INS-evoked PSTHs were classified into onset, sustained, onset-sustained, and offset categories representative of the known range of possible responses(72) (Fig 3A). PSTHs showing post-stimulation drop of 95% of basal activity were assigned an “inhibitory” flag corresponding to presence of post stimulus inhibition. Classification results are summarized in Table 1. Onset responses were the most represented class (Onset+Inhibition: 49.93%, Onset: 12.04%) followed by sustained (Sustained: 18.78%, Sustained+Inhibition:4.51%) and onset-sustained classes (Onset-sustained: 6.05%, Onset-sustained+inhibition: 2.82%). Offset responses were the rarest observed class (5.87%). Observed distributions of firing classes are supported by studies of auditory evoked cortical unit responses(72), suggesting that INS drive naturalistic thalamocortical encodings.

**Figure 3.**
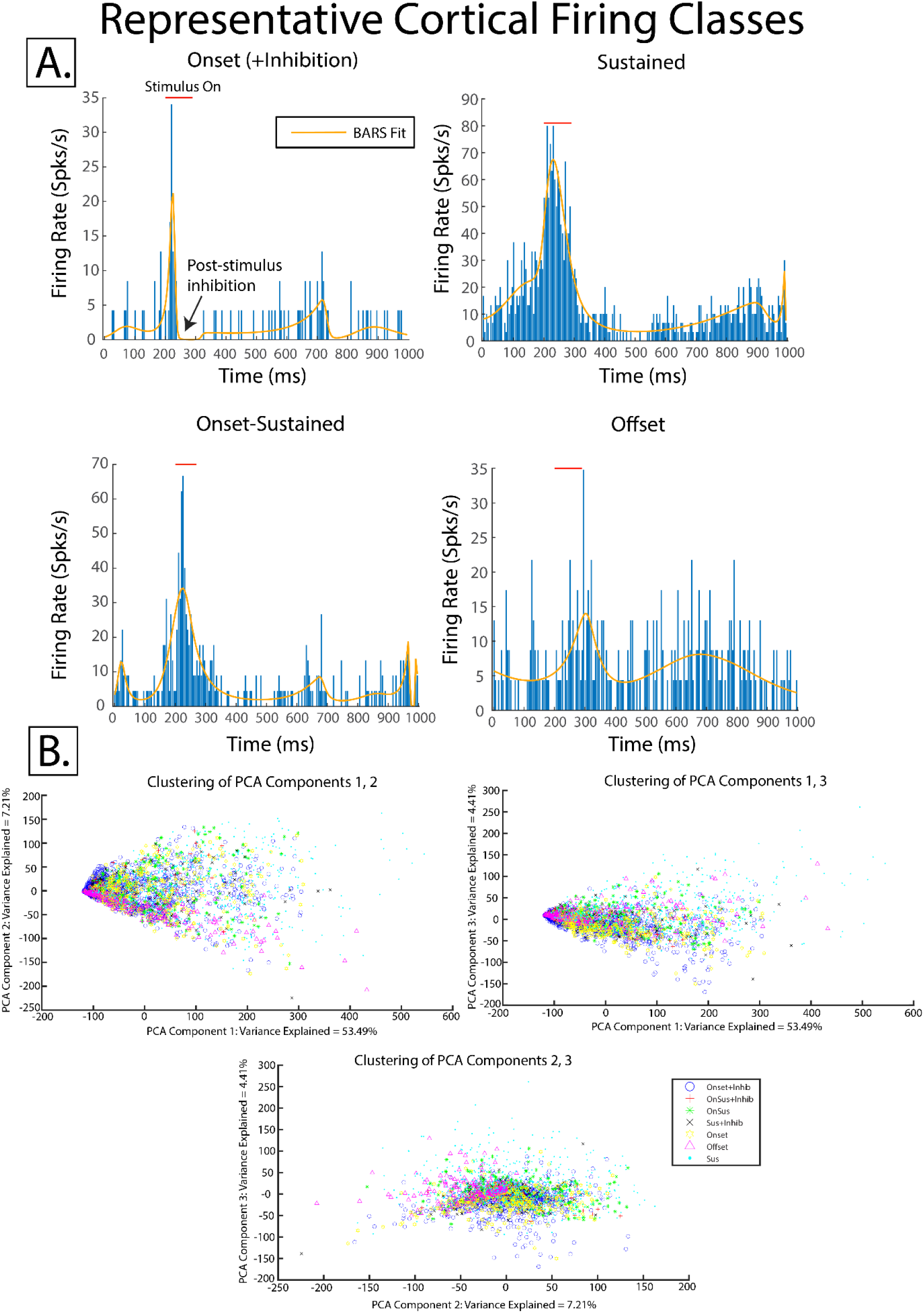
A. Evoked cortical firing activity was classified into onset, onset-sustained, sustained, and offset classes. Any response which showed an offset inhibition resulting in basal firing rate <5% prestimulus firing rate was given an inhibition designation (top left, Onset for example). B. Decomposition of response classes into the top 3 principal components show that these classes exist across a continuum.

**Table 1.** Distribution of Cortical Firing Classes (n=3371)

| **Firing Class** | **% of Responses** | **% of Responses in Class** |
| --- | --- | --- |
| Onset | 12.04 | 61.97 |
| Onset + Inhibition | 49.93 |  |
| Sustained | 18.78 | 23.29 |
| Sustained + Inhibition | 4.51 |  |
| Onset-Sustained | 6.05 | 8.87 |
| Onset-Sustained + Inhibition | 2.82 |  |
| Offset | 5.87 | 5.87 |

While these response states were categorically divided into possible response classes(72), these categories are not meant to suggest all responses fit neatly into well-defined clusters. Principal components analysis (PCA) dimensionality reduction was performed on response profiles to assess the extent to which responses fall on a continuum. Dimensionality reduction into the top 3 components of largest variance (65.11% variance explained) shows that while responses do form some identifiable clusters, responses fall on a continuum of responses within a given cluster, with large overlap between clusters (Fig 3B). Bayesian multinominal regression models (Supplementary Methods) were utilized to infer whether firing class membership was solely a function of INS stimulation parameters. Multinominal regression compares log odds of a PSTH belonging to a given category against a reference category. The most populous onset+inhibition category was chosen as reference. Models suggest that class membership is a function of INS energy and ISI with movement from onset+inhibition to onset resulting from increases in energy and ISI, movement to onset-sustained resulting from decreases in energy and ISI, movement to onset-sustained+inhibition resulting from slight decreases in energy but large decreases in ISI, movement to sustained+inhibition resulting from larger decreases in ISI, and movement to offset class resulting from large decreases in applied energy and smaller decreases in ISI (Fig S12). These models suggest an interplay between INS stimulation parameters, network dynamics, and intrinsic cellular biophysics determines response profile class.

### INS induces spatially constrained thalamocortical recruitment

We next investigated the spatial selectivity of thalamocortical INS using joint peristimulus time histogram (JPSTH) analyses. JPSTHs allow for the assessment of the time-resolved correlation between pairs of neurons in response to INS stimuli. We first assessed stimulation induced correlations of activity related to the initial stimulation event (Fig 4A Left). We next calculated JPSTHs representing functional connectivity between compared neurons when direct stimulus effects are removed (Fig 4A Middle). Consistent spatial geometry of planar recording arrays allowed for assessment of the functional connectivity of responses as a function of distance (Fig 4A right). The maximum spread of correlated activity across all energies was calculated to obtain an upper bound of lateral stimulation spread. Previous electrical mapping studies in rodent auditory thalamocortical areas using linear, Michigan style arrays in nearly all cases showed electrical stimulation spread across the entire extent of recording arrays, up to 1900 *μm* (73, 74). INS correlation analysis shows all responses were constrained to ≤ 1500 *μm*, with 90% of responses constrained to ≤ 1000 *μm* (Fig 4B, left). We next recalculated maximal spread for active units at stimulation intensities < 1mJ, corresponding to an inflection point of increased stimulus transmitted information (Fig 2C), to assess if maximum spatial spread is modulated by INS intensity. At lower energy stimulation, maximal spatial spread was limited to < 1250 *μm*, with 90% of responses constrained to ≤ 1000 *μm* (Fig 4B, right), including numerous instances of moderate correlation even ≤ 500 *μm*. These data suggest maximal spreads of INS-induced activity is significantly less than electrical stimulation. Spread of activation after accounting for direct co-stimulation induced by INS shows similar results, with spreads of correlated activity limited to 1250 *μm* across all energy levels and 1000 *μm* for energies < 1mJ (Fig 4C).

**Figure 4:**
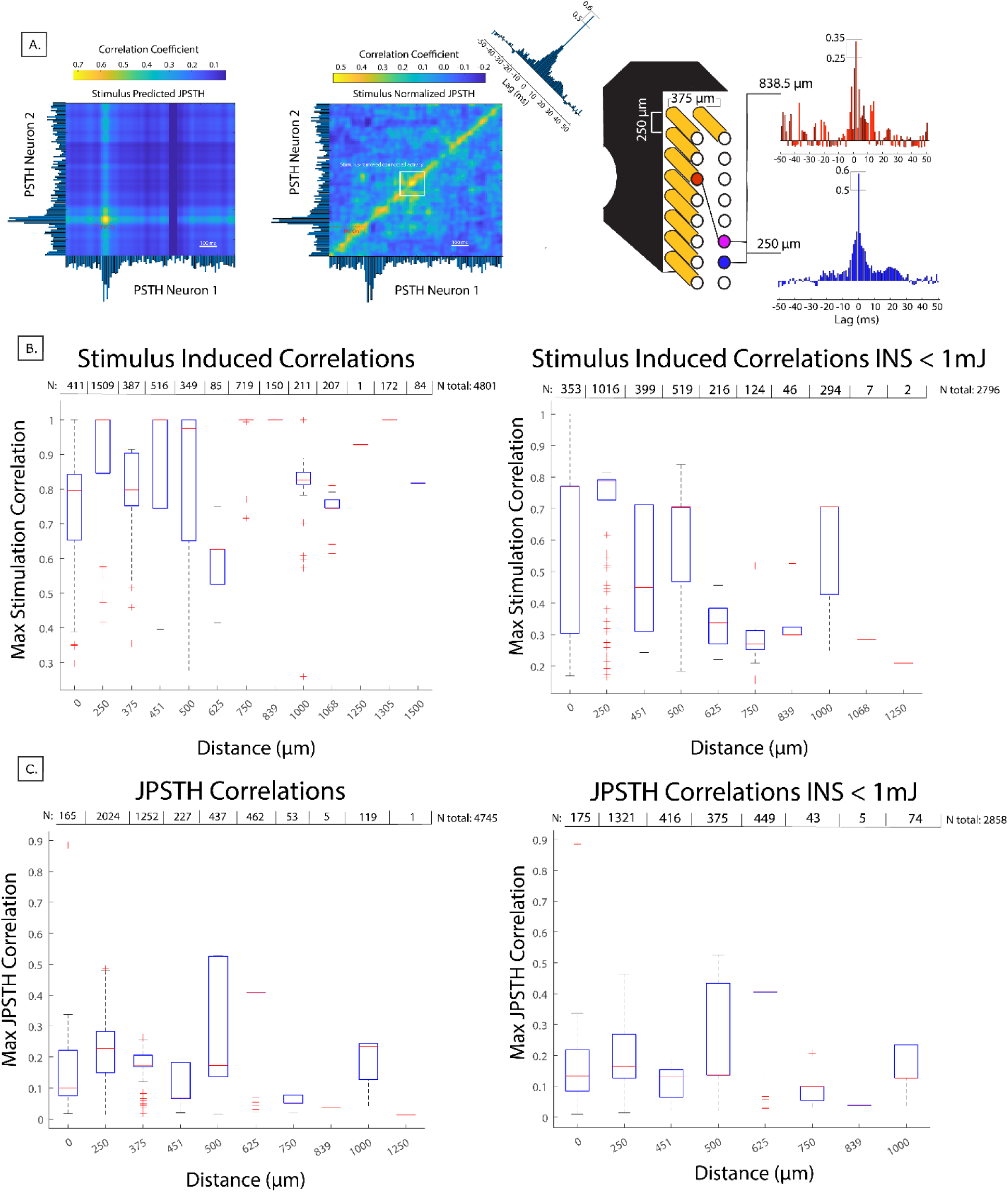
Joint peristimulus time histogram analysis reveals INS thalamocortical recruitment is spatially constrained. A. Schematic of JPSTH analysis. Covariance maps were first calculated between the two PSTHs under test. Covariance maps represent the joint activity of two neurons due to the INS stimulus directly. Subtracting the covariance map from the joint histogram generates the JPSTH, a measurement of correlated activity of the neural network in response to the stimulus. Creating a histogram of the main diagonal of the JPSTH creates a coincidence histogram of total synchrony of the two neurons. Finally, cross correlograms create a statistic of connectivity of the two neurons. Covariance and JPSTH joint histograms were smoothed by a 2D gaussian filter for visualization purposes, but full calculations were performed on raw joint histograms. B. INS-induced correlations show that lateral spread of activation in cortex from thalamic INS were constrained to ≤ 1500 *μm*, with 90% of responses constrained to ≤ 1000 *μm*. Laser energies < 1mJ limited lateral spread to ≤ 1250 *μm*. C. Pairwise JPTHs, measuring post-stimulation induced connectivity show lateral spreads limited to ≤ 1250 *μm* across all applied energies and ≤ 1000 *μm* for stimulus energies < 1 mJ. All correlations and JPSTHs shown were statistically significant (*p* < 0.05) after permutation testing.

### Closed-loop control through deep reinforcement learning

After observation of spatial selectivity in thalamocortical INS, we sought to control small neural populations through closed-loop feedback. Current adaptive DBS systems used in Parkinson’s disease use relatively simple control algorithms centered around reducing β band biomarker correlates of symptomology using single or dual threshold “thermostatic” control(42, 75, 76) which may interfere with activities such as volitional movement(42) and may potentially occlude oscillatory neural dynamics unrelated to disease(39). Control of smaller populations of neurons relevant to disease with control algorithms that encode subject specific firing dynamics may provide targeted treatment and a reduction in off-target side effects. We utilized deep reinforcement learning (RL) to learn complex stimulus-response dynamics in real time while finding stimuli to reach a desired firing states. State, in this study, refers to discrete classes of dynamical activity with stereotyped spontaneous and stimulus evoked activity(77). RL consists of a computational agent which takes actions in response to observations of a given neural state and learns which actions to take to maximize current and future rewards. In our deep RL paradigm, termed SpikerNet, the RL agent can take actions from an action space consisting of INS stimulus parameters of laser energy, ISI, and number of pulses which are constrained to consensus safe energy levels. Stimuli are applied in response to observation of neuron PSTHs from recording electrodes. A reward was then calculated by quantifying mean-squared error distance between evoked firing and target PSTHs. Action policies and state response relationships are then learned using actor-critic deep neural networks, with the actor network encoding actions to take in each environment and the critic learning present and future rewards of taking a given action (Fig 5A, S13).

**Figure 5:**
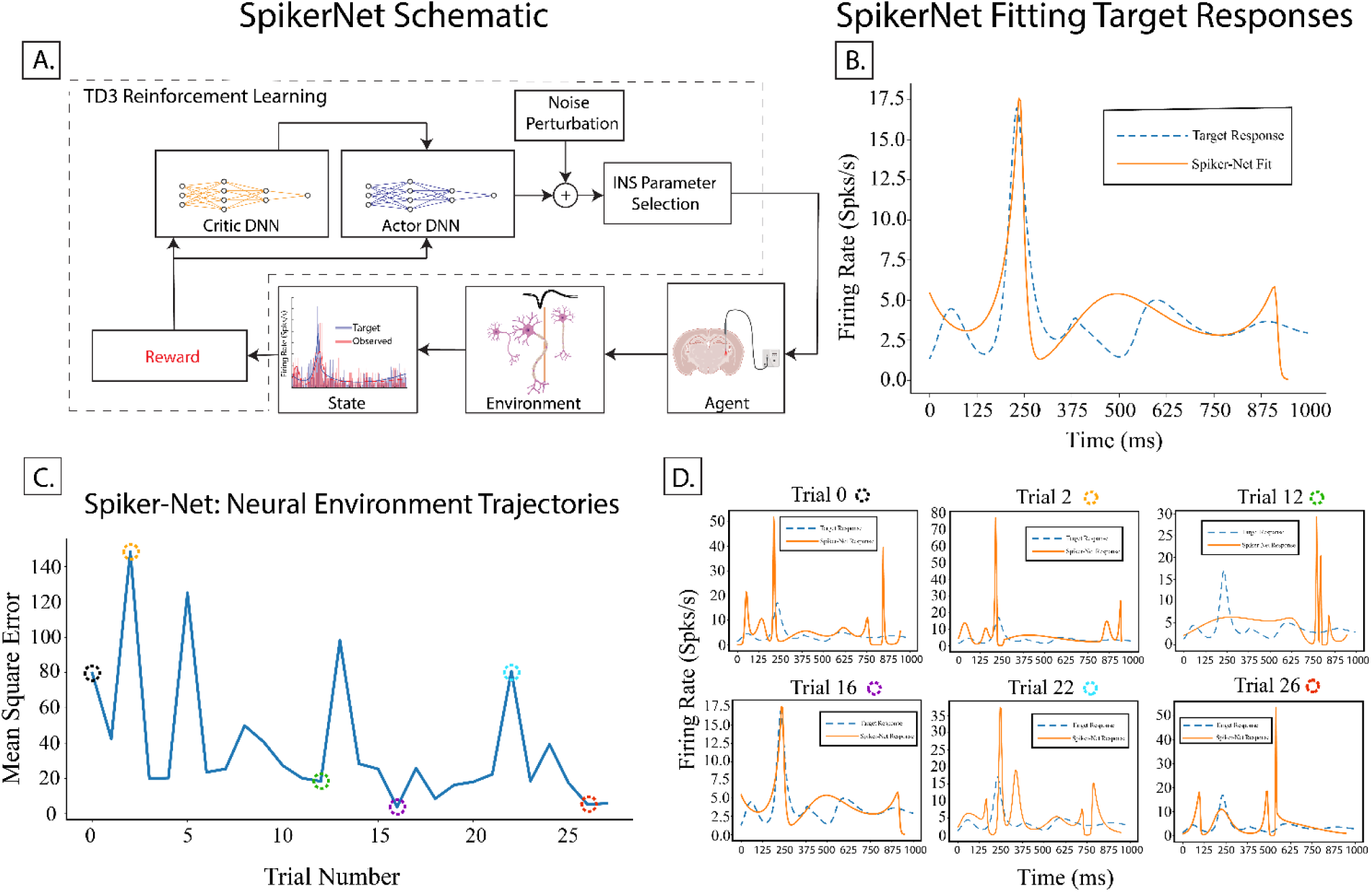
SpikerNet, a deep reinforcement learning based closed loop control system. A. Schematic of SpikerNet operation, which utilizes TD3 reinforcement learning. The state is representative of a response as recorded from the electrode environment. The agent is the set of all safe stimulation parameters. B. SpikerNet is able to find arbitrary neural firing patterns through repeated iterations of stimulation through the environment. C. SpikerNet partakes in search and targeting behavior to find target responses and to learn stimulation parameters which best drive the neural environment to target state. D. Example evoked responses during SpikerNet search and learning show a wide variety of firing classes are evoked during algorithm search. While fits were calculated around the window of evoked activity, more complex multi-peaked and offset responses were observed (Trial 12, 22, 26).

We have previously shown in computational models that SpikerNet is able to quickly learn stimulus trajectories to achieve desired firing patterns(39). SpikerNet’s ability to achieve desired firing patterns *in vivo* was tested by sampling from distributions of previously evoked responses to create novel, previously unobserved firing response target states for the recorded unit. We determined SpikerNet was able to find target firing states precisely (Fig 5B, mean-squared error = 3.872) within a limited number of search iterations (Fig 5C) as predicted in our computational studies(39). It should be noted that search dynamics are intrinsically stochastic and unique to a given animal, target response, and algorithm seeding. Search trajectories during training stages show rapid discovery of target responses indicated by low mean square error followed by exploratory behavior away from the target(Fig S13), characteristic of RL sampling of action-response distributions(78) and necessary to develop a full stimulus to response mapping. We also found that SpikerNet exploration generated a wide variety of firing classes during search that were not identified during our standard intensity and ISI stimulation protocol, including onset-inhibition responses (Fig 5D, trial 0,2), sustained activity followed by burst offset response (Fig 5D, trial 12), and multi-peaked sustained responses (Fig 5D, trial 22). The ability to create and observe such diverse firing patterns is critical to learning stimuli to generate any firing state as well as relearn stimulus-neural dynamics as responses change due to age of recording and stimulation devices and neural adaptation over time.

## Discussion

In this study, we demonstrated INS as a viable oDBS method for treatment of circuitopathy-related neurological diseases and disorders. We quantified INS dose-response profiles and stimulus-response information transformations while also showing the ability of INS to drive biophysically relevant cortical responses at safe energy levels. We further show that INS provides spatially specific activation in thalamocortical networks with spread well below conventional electrical stimulation. Finally, we leverage the spatial specificity of INS to derive a deep reinforcement learning based closed-loop optical control system that can drive neural responses to target states.

### INS drives physiological thalamocortical responses

While many previous INS studies have explored the role of wavelength dependence on INS activation(21, 79, 80), dose-response relationships have largely not been studied. Activation profiles are critical for therapeutic dosing of neuromodulation therapies to titrate efficacious pulses while minimizing patient discomfort from overdriving neurons. Dose-response curves show exponential increases in maximum firing rates in response to increased laser energy with some evidence that extremes in interstimulus intervals further shape neural response PSTHs through integration of INS pulses close in time (Fig 1A,B). One caveat to our study is that only excitatory responses were considered. It has been observed that continuous pulse-width or high frequency (≥ 200 *Hz*) INS stimulation can drive selective inhibitory responses in nerve through introduction of a thermal block(81–83), though this type of stimulation can produce longer lasting mixed excitatory and inhibitory responses, with a higher proportion of excitatory responses for lower stimulus energies(84). An understanding of joint excitatory and inhibitory effects of circuital INS would potentially allow for bidirectional control of local microcircuits and is planned for further study.

INS of thalamocortical neurons produced a variety of short-latency peristimulus responses in auditory cortex neurons, comparable to sound driven auditory cortex responses across species(72, 85–88). These results suggest that thalamocortical INS stimulation largely preserves natural network activation, including sustained responses where the response outlasts the stimulus as well as inhibitory responses. Our INS stimulation parameters differed from previous INS studies in somatosensory cortex and more closely resemble MGB firing rates(51, 89). There is evidence that DBS imparts its therapeutic effect partially through activation of motor cortex from antidromic activation of subthalamic nucleus collaterals(90). While our study can’t strictly rule in or out similar antidromic activation of thalamocortical targets, given that MGV largely sends afferent projections to layer III/IV of A1, INS activation in the present study is likely driven by orthodromic stimulation.

### Spatial selectivity of thalamocortical INS

A oft-touted advantage of INS for cochlear/peripheral (16, 25, 31) and cortical neuron(23) stimulation is constrained stimulation. However, there historically has been a dearth in understanding of network responses and spread of activation through synapses elicited from INS. Previous studies have often focused on intrinsic optical and calcium imaging recording of cortical cells from direct INS stimulation(22, 23, 91). Here we show that INS drives spiking responses across the thalamocortical synapse within a constrained region that is significantly smaller than the region affected by equivalent electrical stimulation. At low INS stimulation energies, activation could be ≤ 500 *μm*, and even at saturating energy levels for firing rates, activation was typically less than 1500 *μm*. It is possible that the activation spread at low energies could be even more restricted, given that we were not able to measure spread of activation in the immediate vicinity of the implanted optrode and that we did not optimize thalamus/cortex overlap in our implantation. Both anatomically and electrophysiologically in A1, there are matched reciprocal projections between the auditory thalamus and cortex(92, 93). Additional mapping during implantation surgery to identify most effective stimulation sites for a given cortical site may reduce energies needed or increase informational capacity even further. As hybrid recording electrodes fixed with optrodes are in use in optogenetic studies, it is feasible to fabricate similar recording arrays with optics that pass near-infrared stimuli, allowing for the study of joint activation and spread in thalamus and cortex concurrently. Regardless, our results show finely graded thalamocortical recruitment, which would potentially reduce off target stimulation side effect profiles in oDBS applications. Further constrained stimulation could also be set during the programming stage of an oDBS system, potentially allowing for fine tuning of therapeutic stimulation.

### Clinical viability of INS

This study lays significant groundwork for the preclinical development of INS for use in a spatially constrained oDBS system. Furthermore, INS has already shown promise in human nerve mapping(31) and intracortical microstimulation(94). However, significant hurdles remain for translation of INS. Laser parameters necessary for stimulation have high optical energy (1-4 mJ) requirements, making fully implantable devices technically challenging. Much progress is currently being made in implantable IR systems that satisfy requirements for stimulation which could be realizable on implantable pulse generators(95). Safety profiles of INS are also promising, with tissue ablation thresholds well studied(29–31, 96, 97). While our data suggests INS drives biophysically relevant responses across a diversity of cell response patterns, disease models are necessary to fully assess therapeutic potential of INS as a DBS paradigm. The biophysical mechanisms of INS are still in debate, with transient thermal gradients(98, 99), transient cellular capacitance changes(100, 101), intracellular calcium cycling(102, 103), intrinsic ion channel light transduction(27, 104), or combinations thereof suggested as causative mechanisms of INS. While not directly assessed within this study, observed short-latency, fast-spiking responses suggest primary ion channel mediation of INS as opposed to slower intracellular calcium signaling. *In vitro* whole cell and outside-out patch clamp studies could elucidate the interplay of the intracellular and membrane bound ion channel sequalae of transient and local thermodynamic changes. A better understanding of these photon-neuron interactions could give rise to more efficient stimulation with larger margins of safety for use in clinical settings.

### Closed-loop reinforcement learning based DBS

Closed-loop DBS provides key advantages over conventional open-loop DBS, including improved stimulation efficacy, reductions in side-effects, and longer IPG battery life(41, 105). However, current closed-loop approaches are limited by non-specific activation of neural targets(76) and relatively simple, threshold-based control algorithms which have difficulty in deciphering pathologic and non-pathologic neural activity(39, 42). We developed SpikerNet to take advantage of spatial selectivity found in INS while also allowing for robust learning of complex neural firing patterns in real-time. An advantage of reinforcement learning over other deep neural network paradigms is that statistical models of neural firing patterns are learned *in situ* and are specific to a subject’s unique neural responses, requiring little training time and not requiring retraining or recalibration. We show that SpikerNet rapidly finds and fits targeted firing patterns (Fig 5B) with search behavior that suggests the ability to fit a wide range of possible neural firing patterns (Fig 5C). We have previously shown in computational models that SpikerNet is flexible to drastic changes in firing patterns(39) suggesting that SpikerNet can adapt to long term changes in neural environments present in chronic, clinical DBS and can reduce the number of trips to the clinic for stimulator adjustments. We also observed evidence of SpikerNet finding target responses through the duration of a subject’s recording period, during which arousal can significantly change firing responses requiring retuning of stimulus parameters (Fig S13). Taken together, SpikerNet could serve as a powerful closed-loop DBS paradigm which can learn and adapt to changes in individual neural responses.

Deep neural network-based approaches however present a significant challenge for translation, in that algorithm decisions are typically made through a “black box” and ultimately unobservable system that may limit guarantees on device efficacy. Reinforcement learning methods however are advantageous in that the stimulus-response relationships after training can be directly observed in implanted devices, allowing for better inference on device operation. However, as stimulation policies are learned using deep neural networks, the salient neural state features leading to stimulus policy formation is still subject to the blackbox problem. The use of novel small network RL policy interpretability tools(106) with *a posteriori* evaluation of trained input/output responses can allow for a deeper understanding of algorithmic decision making. In this way, we see SpikerNet as a tool which can be utilized as a “physician in the loop” system, where SpikerNet can be utilized in concert with a trained DBS technologist to assist in difficulties found in DBS programming(107) and with physician monitoring during autonomous learning and stimulation.

## Materials and Methods

All experimental and surgical procedures and protocols were approved by the Institutional Animal Care and Use Committee (PACUC) of Purdue University (West Lafayette, IN, #120400631) and in accordance with the guidelines of the American Association for Laboratory Animal Science (AALAS) and the National Institutes of Health guidelines for animal research. A total of 11 rats were used in this study.

### Surgical Procedures

Adult Sprague-Dawley rats with weights between 300-400 g (Envigo, Indianapolis IN) were initially anesthetized in an induction chamber with 5% isoflurane and given a bolus injection of a ketamine/dexmedetomidine cocktail (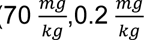 respectively). Surgical plane of anesthesia was monitored continuously throughout the procedure by evaluation of toe-pinch reflex. A preoperative analgesic dose of Buprenorphine (1 mg/kg) was administered 30 minutes prior to first incision and every 6-12 hours for 72 hours post-surgery. Rats were placed in a stereotaxic frame secured by hollow ear bars. An initial incision was made down midline with blunt dissection of periosteum performed to reveal cranial sutures. Three stainless steel bone screws were placed in the skull to ensure stability of implanted devices and headcap with a fourth titanium bone screw placed to serve as a ground and reference electrode(108). Right hemisphere temporalis muscle was gently resected and a 2×2 mm craniectomy was made above auditory cortex (A1) (centered: -6 AP, -5 ML)(109). Dura was gently resected using a 25G curved needle. A 2mmx2mm 16 channel microwire array (TDT, Alachua FL, electrode spacing given in Fig 1A) was inserted perpendicular to the surface of the brain. Devices were slowly inserted into A1 during application of 80 dB gaussian noise stimuli. Devices were placed centered putatively in layer III/IV of A1 after confirmation of low latency, high amplitude multiunit activity was observed on the array(49, 110). One animal received a 3mm linear array (NeuroNexus A1-16, 200 *μm* between contacts) with contacts placed in A1 layers 3/4 in place of TDT planar array. A second craniectomy was made above the medial geniculate body (MGB) (-6 AP, -3.5 ML)(49) and a fiber optrode (Thor Labs, Newton NJ) was placed -6 mm into tissue (Fig 1A). Recording arrays and fiber optics were sealed into place by application of UV-curable composite (Pentron, Wallingford, CT). Rats were returned to their home cage and allowed to recover for 72 hours prior to beginning of the recording regime.

### Electrophysiological Recordings

All recordings were performed in a 9’x9’ electrically and acoustically isolated chamber (Industrial Acoustics Corporation, Naperville IL) with laser electronics placed outside of the chamber to prevent field interactions from high current pulses(111, 112). Prior to recording sessions, rats were given a bolus intramuscular injection of dexmedetomidine (0.2 mg/kg) for sedation(49, 53, 113). Optical stimuli were delivered via a custom made, open-source INSight system (all plans available at our Github repository: https://github.com/bscoventry/INSight and included in supplementary material) with a 1907 nm semiconductor laser (Akela Trio, Jamesburg NJ) fiber coupled to the optrode with a 200 μm, 0.22 NA fiber (Thor Labs FG200LCC). Laser stimuli were controlled via a RX-7 stimulator (TDT) and consisted of train stimuli with pulse widths between 0.2-10 ms, interstimulus intervals between 0.2-100 ms and energy per pulse between 0-4 mJ, below reported thresholds of laser ablation(28, 31).

Each recording trial was composed of a 200 ms pre-stimulus interval to facilitate spontaneous rate calculations, application of the train stimuli, and a post-stimulus interval with total trial length equal to 1 second. Applied laser energies were randomized to limit effects from neural adaptation with 30-60 repetitions per pulse width/interstimulus interval combinations. Signals from recording electrodes were amplified via a Medusa 32 channel preamplifier and discretized and sampled at 24.414 kHz with a RZ-2 biosignal processor and visualized using Open-Ex software (TDT). Action potentials were extracted from raw waveforms via real-time digital band-pass filtering with cutoff frequencies of 300-5000 Hz, with LFPs extracted from real-time digital filters with bands 3-500 Hz. Chronic recordings were made through the lifetime of implanted optrodes and electrodes. To assess the impact of unilaterally implanted devices, pre and post-surgery mid-latency response (MLR) electroencephalography was performed. The experimental setup used has been described in detail in previous studies(54, 55) and further explained in supplementary methods.

### Data Processing and Analyses

Action potentials and MLRs were exported and processed using custom written programs in the Matlab programming environment (Mathworks, Natick MA). Spikes were sorted into single-units using superparamagnetic clustering methods in Wave-Clus(114). Peri-stimulus time histograms (PSTH) were calculated and density estimates of firing rate functions were calculated from PSTHs using Bayesian adaptive regressive splines (BARS) under a Poisson prior with *λ* = 6 (115, 116). Trials containing artifacts due to breathing or volitional movement were detected via between-channel cross correlation and RMS voltages exceeding 1 mV were removed from recordings. To facilitate comparisons between electrodes and animals, PSTHs were standardized using the following equation:

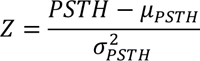

where Z is the standardized PSTH and 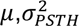 are the mean and standard deviation of the PSTH. Neurons were classified as responsive to INS if a PSTH in the stimulus series showed a z-score firing increase of ≥ 7.84 (4 ∗ 1.96, 1.96 = critical Z-score threshold) above mean spontaneous firing rate.

After detection and PSTH calculation, single unit responses were sorted into one of 7 established firing pattern classes found in rat auditory cortex(72, 117). Responses were first classified into onset, offset, sustained, or onset-sustained classes, with onset responses exhibiting a rise above spontaneous activity followed by a drop to spontaneous rates before cessation of the stimulation and offset responses characterized by an increase in firing rate from baseline after termination of stimulus plus 7ms to account for maximal response latencies in cortex from thalamic stimulation(117, 118). Responses showing firing activity above spontaneous activity throughout the duration of the stimulus were classified as sustained or onset-sustained, with onset-sustained responses showing a ratio of peak onset response to sustained rates >3. The inhibited response subclass showed a post-stimulus reduction in basal firing rate to below 95% of mean rate during the 200 ms prestimulus interval.

Mutual information (MI) measures of thalamocortical encoding of INS stimulation were calculated using the methods of Borst and Theunissen(114) with bias correction performed using quadratic extrapolation(63). Full information theoretic calculations are provided in the supplementary methods. Lateral spreads of activation across cortical neurons were assess through joint peristimulus time histogram (JPSTH) analyses(119). Full JPSTH models and algorithmic descriptions are given in the supplementary methods.

### Deep Reinforcement Learning Based Closed Loop Control

Closed loop DBS control was achieved through a novel deep reinforcement learning based paradigm which we termed as SpikerNet(38). SpikerNet was programmed in Python using the Pytorch deep learning backend(120). A custom made OpenAI Gym environment served as the interface between TDT data acquisition hardware and Pytorch. Deep reinforcement learning seeks to maximize a target reward by continually sampling an environment while learning which actions taken provide highest future rewards through time(121, 122). In SpikerNet, the environment space was defined as the continuum of evoked cortical neuron firing rate PSTH densities. The action space as the continuum of stimulation amplitudes, pulse widths, and number of INS pulses delivered in a trial. The action space of stimulation parameters was limited in both hardware and software to below ablation thresholds to ensure SpikerNet did not damage thalamic structures during parameter search. Deep reinforcement learning was performed using the twin-delayed deep deterministic policy gradients (TD3) algorithm, which is a model agnostic double Q learning method for continuous environment and action spaces that outperforms other model-free deep-Q learning methods(123). To assess the ability of SpikerNet to reach arbitrary spike PSTHs, distributions of all observed PSTHs were formed. From that distribution, a target PSTH was sampled and represented a non-observed but biophysically plausible target PSTH. Reward functions were set as

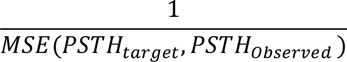

with mean-squared error (MSE) chosen as it provides asymptotically the maximum likelihood estimator. Online multi-unit PSTHs were calculated online from 10 repetitions of INS stimuli with densities estimated using online Bayesian adaptive regression splines. A MSE value below 0.14 denoted an observed result that is sufficiently close to the target response and acts as a signal to begin a new search episode. It is important to note that SpikerNet performs reward maximization through all episodes and is not truncated at the threshold of a sufficiently close fit.

### Statistical Methods

All statistical methods, models, and sensitivity analyses are given in supplementary methods and materials.

## Acknowledgments

This study was supported by a grant from the National Institutes of Health (NIDCD R01DC011580, PI: ELB) and the Purdue Institute for Integrative Neuroscience collaborative training grant (PI: BSC). The authors would like to thank Dan Pederson, PhD and Kirk Foster for guidance in PCB layout and Don Ready, PhD for maintenance and funding of the Purdue University Life Science Fluorescence Imaging Facility. B. S. Coventry is now affiliated with the Department of Neurological Surgery and the Wisconsin Institute for Translational Neuroengineering, University of Wisconsin-Madison, Madison, WI USA.

## Data and Code Availability

All neural analysis and statistical inference code is available from https://github.com/bscoventry/OpticalTCNeuromodulation. Data used in this study can be found in the following open science framework repository: https://osf.io/w4ufh/. INSight INS system build files and materials list is available from: https://github.com/bscoventry/INSight. Due to patent restrictions, data and source code related to SpikerNet is available upon reasonable request from corresponding authors.

## Supporting Information

### 1. Bayesian Model Descriptions and Sensitivity Analyses

This report follows the guidelines for reporting of Bayesian Analysis (BARG) (1) consisting of:

- Necessary software and source code directory
- Goals of the analysis
- Model descriptions and decision criterion
- Prior and hyperprior descriptions
- Sensitivity analyses for varying prior distributions
- Posterior and MCMC diagnostics

#### 1.1 Necessary software and source code directory

BARG: Step 2A, 6

Bayesian modeling was performed using Python 3.6.8 on an MSI GS-66 Laptop with an Intel Core i7 processor (6 cores) and an Nvidia RTX2070 GPU. Models were implemented in PyMC3 version 3.11.5 (2), a probabilistic programming module in the Python environment. All source code is available at this paper’s github repository (SI: Software S1). All source data is available at this article’s open science framework repository (SI: Dataset D1).

#### 1.2 Goals of the Analyses

BARG: Preamble

The goal of the utilized regression analyses is to establish a model of the relationship between stimulus parameters (applied laser energy and interstimulus intervals) with evoked thalamocortical neural responses as quantified by single unit firing rates. While this is normally established using frequentist multilinear regression analyses, neuron responses are heterogeneous with differences arising from nominal firing patterns arising individual cell types, differences in exact placement in receptive fields of stimulation and recording devices, and changes in within-animal recordings from glial scaring and skull growth over time leading to changes in placement of devices. These nuances are best characterized by hierarchical regression models.

Bayesian approaches allow for flexible and explicit hierarchical model descriptions which provide rich and descriptive inference and quantification of uncertainty in measurements by inference of direct probability measures on posterior distributions as opposed to less intuitive and harder to interpret frequentist p-values. Bayesian approaches are data driven and account for previous knowledge to be encoded as prior distributions. It can be shown that Bayesian hierarchical regression is a regularized frequentist random effects model with uniform distributions on the hyperparameters. However, frequentist approaches collapse inference into singular decision boundaries (p-values) and do not allow for model constructions which best fit the observed data.

To this end, we utilized Bayesian hierarchical multilinear regression to account for both within and between subject differences of evoked responses to INS stimuli as a function of applied laser energy, time between laser pulses (interstimulus intervals, ISI), and the interaction between applied laser energy and ISI. The general regression model is:

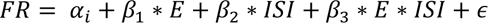

where FR is the max evoked firing rate. Firing rate functions were calculated from recorded peristimulus time histograms with Bayesian adaptive regression splines density estimation(3). Parameters *β*_*i*_ quantify the effect of laser energy(*β*_1_), pulse ISI(*β*_2_), and laser energy and pulse ISI interaction(*β*_3_) on evoked firing rates respectively. The *α* parameter describes the model intercept and quantifies subthreshold spontaneous activity and the *ϵ* quantifies model error.

Hierarchical models perform ‘partial pooling’ of response data which accounts for individual differences in parameter estimation. This is done by assuming parameters *α*, *β*_1_, *β*_2_, *β*_3_ and *ϵ* are not singular values but form distributions that quantify firing rate dependencies from laser parameters while accounting for within-subject differences. Prior selection is discussed in section 1.3.

Partial pooling was performed by adding an implicit class definition *e*_*i,j*_ in PyMC (see code) which encodes the response arising from the *i*^*th*^ electrode in the *j*^*th*^ subject.

We also utilized Bayesian formulations of multinominal regression (aka Softmax regression) to understand the dependency of neural firing class on the laser parameters of applied energy and interstimulus interval. The multinominal regression model is as follows:

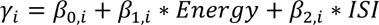

where *γ*_*i*_ is defined as the *i*^*th*^ firing class. Firing classes included onset, onset-sustained, sustained, onset+inhibition, onset-sustained+inhibition, sustained+inhibition, and offset with class inclusion criteria defined in the materials and methods portion of the main text. The probability of class *γ*_*i*_ is calculated via the softmax link function:

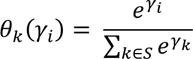

which creates a mapping of outcome *γ*_*i*_ against all *k* outcomes in the set of possible classes *S*. For efficient computation, *β*_1_ and *β*_2_ were recast as a singular tensor optimized for sampling. The multinominal regression model then takes the following form:

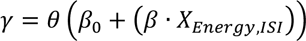

where the · operator is the tensor inner product.

#### 1.3 Prior Selection

As single unit thalamocortical recordings elicited from INS have largely been unexplored, previous knowledge cannot be adequately constructed into a highly informative prior. However, our previous experience in thalamus and cortical recordings(4–10), the central auditory pathway (11–13), and in INS parameter selection(14, 15) gives prior information on potential variances of firing rate in cortex from thalamic stimulation. As such, we chose moderately informative distributions (see section 3) so as to not unduly influence the posterior and let the observed data fully inform the posterior. Normal distributions were chosen over uniform distributions to allow for unforeseen high variance, low probability events to inform the posterior if evidence is sufficiently strong. A choice of uniform distribution would drive such events to probability zero, missing potentially notable neural recruitment. To ensure the prior distribution did not unduly inform posterior distributions away from observed data, sensitivity analyses to prior parameters was performed (Section 3).

Observation of evoked INS responses tended towards normal distributions, dictating a normal likelihood distribution. Previous studies in regression suggest the use of a Student T distribution, which incorporates an added hyperprior for degrees of freedom (*v*), performs a robust regression against potential outliers(16). Importantly, as *v* → ∞, the Student T distribution becomes a normal distribution and relative tail spread of the Student T distribution is learned online through *v* hyperpriors.

An interaction term, *β*_3_ ∗ *E* ∗ *ISI* was included in the analysis as it was hypothesized that extremely short ISIs could cause neuron interactions between pulses potentially leading to temporal integration of laser energy.

#### 1.4 Posterior Decision Rules

Inference was performed on posterior distributions with credible regions (analogous to frequentist confidence intervals) defined as a highest density interval (HDI) of 95% of parameter maximal *a posteriori* density (MAP) parameter estimates which represent the most probable value of the coefficient. MAP estimates are analogous to maximum likelihood estimation found in frequentist approaches. This allows for the quantification of parameter uncertainty as variance observed in posterior parameter distributions, with narrow HDIs representing more certain estimates. It is customary to define a region of practical equivalence (ROPE) if prior information dictates that incremental parameter changes are effectively the same. As we lack prior knowledge to inform the choice of a prior rope, we take an agnostic approach that any change seen is worth investigating and thus ROPEs are not presented. An effect was deemed significant if it’s 95% HDI did not overlap with 0, in line with proposed decision rules typical of Bayesian inference(17, 18).

#### 1.5 Final Model

Posterior predictive checks and sensitivity analysis were performed to titrate the best performing models as measured against observed data (Section 3). The final hierarchical regression model is schematized in figure S1 and for the multinominal regression model in figure S2. Final models included deterministic nodes at outputs of prior nodes to prevent NUTS from becoming stuck in regions of the sampling space which are difficult to explore ^1^.

#### 1.6 Model Sensitivity analyses

BARG: Step 3A,C

Individual To evaluate the dependance of hyperprior parameters on model fitting, we used leave one out (LOO) cross validation(19). Three separate models were evaluated with model variances varied to test sensitivity of each model. Initial model construction suggested that natural-log transformations of the dependent variable (firing rate) produced distributions which are better modeled as normal distributions. To this end, hierarchical models under test were as follows:

**Table S1:** Regression models under test.

| MODEL NAME | MODEL |
| --- | --- |
| REGRESSION | $FR = \alpha + \beta_1 * Energy + \beta_2 * ISI + \beta_3 * Energy * ISI + \epsilon$ |
| SEMILOG REGRESSION | $\ln(FR) = \alpha + \beta_1 * Energy + \beta_2 * ISI + \beta_3 * Energy * ISI + \epsilon$ |
| NATURAL LOG REGRESSION | $\ln(FR) = \alpha + \beta_1 * \ln(Energy) + \beta_2 * \ln(ISI) + \beta_3 * \ln(Energy) * \ln(ISI) + \epsilon$ |

For each model, the variance hyperprior was varied to assess the impact of prior parameters on posterior predictions. Prior classes were defined as: informative (variance ≤ 1), moderately informative (variance = 5), and weakly informative (variance ≥ 10). Primary metrics for model comparison were expected log pointwise predictive density (ELPD), defined as(20):

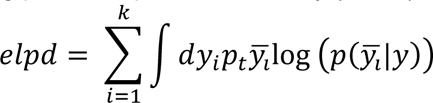

where *p*_*t*_, *y*_*i*_ are unknown distributions representing the true data generating function for estimates of true posterior predictive function 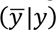 from observed data y. Estimated *p*_*t*_, *y*_*i*_ distributions are obtained via cross validation during LOO analysis. In general, higher values of ELPD are a result of higher out of sample predictive fit indicative of a better model. Weight values generated by LOO cross validation were also analyzed and predict the probability of each model given observed data. Finally, we observed the standard error of the ELPD estimate (SE), and the difference between the model with highest ELPD and every other model (dSE) with dSE of the top model set to 0.00 by definition. All LOO calculations were performed *post hoc* with the python package arviz, a plugin for PyMC.

**Table S2:** LOO model comparison results for the Bayesian hierarchical regression models. Var: Prior variance parameter, log: log predictor and predicted variable model. semilog: semilog predictor model. ST: Student T Likelihood models. N: Normal likelihood models

| MODEL | R | ELPD | WEIGHT | SE | DSE |
| --- | --- | --- | --- | --- | --- |
| **ST LOG VAR 5** | 1 | -5337.48 | 2.046623e-01 | 46.220458 | 0.00 |
| **ST LOG VAR 100** | 2 | -5337.62 | 1.763745e-01 | 46.227682 | 0.420867 |
| **ST LOG VAR 0.5** | 3 | -5337.76 | 1.552051e-01 | 49.173347 | 0.409773 |
| **ST LOG VAR 25** | 4 | -5338.15 | 1.058393e-01 | 46.358847 | 0.492297 |
| **ST LOG VAR 10** | 5 | -5338.18 | 9.996540e-02 | 46.141502 | 0.300197 |
| **ST LOG VAR 1** | 6 | -5338.26 | 9.238175e-02 | 49.030330 | 0.331152 |
| **N LOG VAR 10** | 7 | -5340.60 | 7.103823e-02 | 49.024680 | 3.308668 |
| **N LOG VAR 1** | 8 | -5341.09 | 4.291607e-02 | 48.985814 | 3.293779 |
| **N LOG VAR 5** | 9 | -5341.16 | 3.978488e-02 | 89.737613 | 3.296273 |
| **N LOG VAR 0.5** | 10 | -5342.46 | 1.183257e-02 | 89.930943 | 3.300550 |
| **ST SEMILOG VAR 1** | 11 | -5466.76 | 4.359604e-37 | 84.933022 | 15.845916 |
| **ST SEMILOG VAR 5** | 12 | -5467.12 | 3.535240e-37 | 89.043113 | 15.856552 |
| **ST SEMILOG VAR 10** | 13 | -5467.15 | 5.622764e-37 | 85.266895 | 15.895646 |
| **ST SEMILOG VAR 0.5** | 14 | -5467.18 | 3.483572e-37 | 85.266895 | 15.866405 |
| **ST VAR 1** | 15 | -15336.31 | 0.000000e+00 | 49.465018 | 79.406629 |
| **ST VAR 0.5** | 16 | -15355.67 | 0.000000e+00 | 49.509352 | 80.415787 |
| **ST VAR 5** | 17 | -15355.67 | 0.000000e+00 | 49.487001 | 80.415787 |
| **N VAR 10** | 18 | -16119.11 | 0.000000e+00 | 49.510419 | 82.384329 |
| **N VAR 1** | 19 | -16132.23 | 0.000000e+00 | 49.524316 | 83.549811 |
| **N VAR 0.5** | 20 | -16154.55 | 0.000000e+00 | 49.514661 | 84.262219 |

Sensitivity analyses were also performed for the Bayesian multinominal regression models, also with informative (variance = 1), moderately informative (variance = 5) and weakly informative (variance = 10) prior parameters. Table S3 outlines LOO model comparisons for the Bayesian multinominal regression model. Comparisons suggest that the standard models perform better than semilog based models, with only minimal changes in model performance with varying prior variances, suggesting that the posterior distribution is largely driven by observed data.

**Table S3:** LOO model comparison results for the Bayesian multinominal regression models. Var: Prior variance parameter, semilog: semilog predictor model.

| MODEL | R | ELPD | WEIGHT | SE | DSE |
| --- | --- | --- | --- | --- | --- |
| VAR 1 | 1 | -3861.38 | 0.400151 | 38.15 | 0.00 |
| VAR 10 | 2 | -3861.76 | 0.261702 | 38.79 | 0.84 |
| VAR 5 | 3 | -3861.85 | 0.239520 | 38.82 | 0.88 |
| SEMILOG<br>VAR 1 | 4 | -3871.25 | 0.037136 | 37.59 | 7.28 |
| SEMILOG<br>VAR 10 | 5 | -3871.38 | 0.032181 | 38.12 | 7.36 |
| SEMILOG<br>VAR 5 | 6 | -3871.46 | 0.029310 | 38.12 | 7.36 |

#### 1.7 Posterior and MCMC Diagnostics

BARG: Step 1E, 2A-D, 3A,C

##### 1.7.1 Choice of MCMC method

For sampling, the Hamiltonian-based MCMC method no U-turn sampling (NUTS)**(21)** was used. NUTS presents a modification of general Hamiltonian Monte Carlo samplers and presents an efficient sampler for hierarchical and high-dimensional models at the cost of slower sampling times. Hierarchical regression models ran 4 simultaneous chains with 4000 burn in samples and 5000 iterations with a 95% target inclusion probability. Multinominal regression models also ran 4 simultaneous chains with 5000 burn in samples and 5000 iterations with a target inclusion probability of 99.95% inclusion probability owing to a more difficult posterior to sample.

###### MCMC Diagnostics

Energy transition plots were used to assess how well NUTS explored the target posterior distribution of the best performing model as assessed by PSIS-LOO comparisons between models**(22)**. As NUTS sampling is based off dynamical systems modeling (Hamiltonian Monte Carlo), movement through the typical set towards a target distribution has associated momentum and thus potential and kinetic energy associated with movement through probability space. Efficiency in movement through the target distribution can then be assessed by comparing energy associated with the marginal energy distribution, quantifying the geometry of the underlying target distribution with the energy associated with the distribution of Markov state transitions. The hierarchical regression model displayed overlapping marginal energy and energy transition distributions (Fig S3) suggesting that sample to sample movement was nearly independent and indicative of efficient sampling of the target posterior distribution.

Furthermore, traces of sampled prior and hyper-prior parameters in hierarchical and multinominal regression models suggest effective sampling of the posterior distribution (Fig S4). Furthermore, the Gelman-Rubin statistic, quantifying within and between chain estimates and correlation was 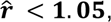 indicative of convergence of marginal posterior parameter values**(23)**.

##### 1.7.2 Posterior Predictive Checks

An advantage of using Bayesian-based inference approaches is the ability to directly and explicitly compare model fits to observed data, a process often not available in frequentist-based software packages or left out of final analysis. During inference model development, posterior predictive checks were performed by sampling from the posterior distribution (20,000 draws). Kernel density estimates of posterior predictive distributions were compared to kernel densities of observed data. Goodness of fit was quantified using the Bayesian formulation of the p-value**(16)**. Similar to the frequentist p-value, the Bayesian p-value is also a measure of discrepancy, quantifying the probability that posterior predictive-based draws are more extreme than observed data. The Bayesian p-value is defined as:

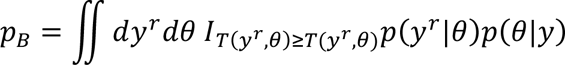

where I is the indicator function, *y*^*r*^ is the posterior predictive distribution and y is the posterior distribution. Similar to the posterior distribution, posterior predictive distribution and Bayesian p-values were estimated using NUTS. The closer the Bayesian p-value is to 0.5, the better the data sampled from the posterior distribute around the observed data.

Figures S5 and S6 display posterior predictive fits and Bayesian p-values for the hierarchical linear and multinominal regression models respectively. Both models suggest excellent posterior predictive fits with 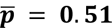 for the hierarchical linear regression model (Fig S5) and 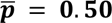 for the multinominal regression model (Fig S6).

#### 1.8 Prior and Posterior Trace plots

BARG: Step 2B,C

Critical to the performance of HMC based Bayesian sampling is the convergence of sampling traces. Output trace plots display the chain of sampled values and the resulting kernel density estimates of sampled distributions. All sampled traces showed no divergences in sampling, suggesting that sampled traces were well behaved in sampling the space of the distribution. Furthermore, the Gelman-Rubin statistic, quantifying within and between chain estimates and correlation was 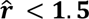 for all sampled traces suggesting good convergence and effective sampling of target distributions. For clarity and transparency, traces are available on open science framework, with traces for all hyperpriors and posteriors presented here (Figures S7-S9). Traces were checked for characteristic sampling behavior**(21)** with no pathological traces found in models.

### 2. Design of the INSight System

#### 2.1 System Description

The goal of INSight was to develop an low cost INS system that was in reach for neuroscience and neuroengineering laboratories who don’t already have access significant optical. To this end, designs were made using off the shelf components with a stimulation interface which could easily connect to recording hardware with DACs or digital trigger outputs.

The INS lasing system consists of 3 subsystem realizations (Figure S10 A); the laser diode, laser diode driver, and driver control system, with two optional but recommended support systems: the thermal electric cooler (TEC) system and the power monitoring system. The laser diode system consists of an Akela Laser (Jamesburg, NJ) Trio diode mounted via pogo pins to a custom interface printed circuit board (Figure S11). The Trio diodes have small form factors, low cost, and are modular, allowing for the rapid switching between different wavelength modules to vary optical penetration depths(24, 25) or to leverage the same system for optogenetic applications. For this study, a 1907 nm, 1W diode was used in accordance with commonly used wavelengths in INS(26). The laser diode is driven through high current, single strand 18-gauge wire using a PLD1000 pulser (Wavelength Electronics, Bozeman MT) with output current controllable by voltage pulses from a control source such as neural recording hardware or the Arduino control module provided. Pulse shapes and waveforms can be implemented using a computer controlled arbitrary function generator connected to the driver analog input. In this study, a RX-7 stimulation isolator (Tucker-Davis Technologies, Alachua FL) was used. In settings where a triggerable voltage source is unavailable, we developed an open source, Arduino based stimulation system which is included in our online build materials. Power to the system was provided by a high current fixed voltage PCB mount supply (LS50-5, TDK-Lambda). The diode laser was fiber coupled to implanted devices using an SMA to FC/PC 400-2400 nm wavelength multimode fiber (Thor Labs, Newton NJ) which can be commutated using a rotary patch cable (Thor Labs). Laser output power was measured using a S305C power sensor (Thor Labs) to validate applied energy. The akela trio substrate displayed strong linearity through periodic open loop tests over the span of a year (Figure S10.B). It should be noted that if the system is operated in open loop, routine power calibrations should be performed to account for laser power drift due to age of the device, ambient temperature, or recent thermal contraction/expansion of the laser substrate.

Modularity was designed into INSight, making it suitable for INS, optogenetics, and as a laser activation source for calcium imaging and other optical techniques. Furthermore, the trio module can be substituted for multiwavelength modules for more expansive applications. INSight can also be modified to accommodate other commercially available laser diode modules.

#### 2.2 Laser Board Layout and Interfacing

In order to create an interface allowing for quick laser diode substitutions, a customized printed circuit board (Figure S11) was created. The pads corresponding to the laser diode should be populated with pogo pins (Adafruit Industries, New York, NY). The laser is place on the pogo pins and secured by screwing the laser diode into the board via through holes on the Trio enclosure, facilitating a quick and easy laser diode replacement system. Traces between the diode laser anode and cathode and laser driver should be as wide as possible with a minimal connection path to ensure proper current handling with minimized trace heating. If wires are used, wire of gauges 18 or less or high current capacity wire should be used with the shortest wire length possible.

#### 2.3 INSight electrical properties

**Table S.4:** Laser System Electrical Properties.

| *Parameter* | *Value* | *Unit* |
| --- | --- | --- |
| *Maximum Laser Output Power* | 1 | W |
| *Laser Diode Forward Voltage* | < 2.5 | V |
| *Laser Driver Analog Input Rise Time* | 7 | μs |
| *Max Laser Driver Supply Voltage* | 5.5 | V |
| *Laser Driver Analog Input Maximum Voltage* | Supply Voltage + 0.5 | V |
| *Laser Driver Current to Voltage Transfer Function* | 4.6 | A / V |

Relevant electrical properties of the proposed system are found in table S.4.

### 3. Supplementary Methods

#### 3.1 Mid-Latency Response Electroencephalography

To assess the impact of unilaterally implanted devices, pre and post-surgery mid-latency response (MLR) electroencephalography was performed. The experimental setup used has been described in detail in previous studies(8, 27).

Briefly, recordings were performed in a double-walled acoustically isolated anechoic chamber. Rats were given a bolus injection of dexmedetomidine (0.2 mg/kg) and maintained at 37℃ via a warming pad. Needle electrodes (AMBU, Columbia MD) were placed in a four-channel configuration (Fig. 2) (channel 1 - Fz to Cz, channel 2 - horizontally P5-P6, channel 3 - contralateral to speaker, C3-P5, channel 4 - ipsilateral to speaker, C4-P6). The reference electrode was placed across the mastoid bone, and the ground electrode placed at the base of the tail. Auditory click stimuli consisting of square pulses of alternating polarity with 0.1 ms in duration at a presentation rate of 4 Hz with sound levels between 65-85 dBSPL. 200 repetitions were collected over a 100ms window and averaged. Presurgical recordings were performed 24-48 hours before surgical procedure and postsurgical recordings were performed 72-96 hours post-surgery.

#### 3.2 Information Theoretic Calculations

Mutual information (MI) measures of thalamocortical encoding of INS stimulation were performed using direct estimation of response distributions from observed data followed by sampling bias correction using quadratic extrapolation(28). Stimulus-information relationships were estimated using the approach of Borst and Theunissen(29):

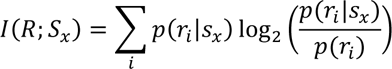

where *I*(*R*; *S*_*x*_) is the “plug-in” estimated mutual information of response *R* conditioned on INS stimulus with stimulus energy *x* (*S*_*x*_) across the duration of the entire stimulus. The values (*r*_*i*_) and *p*(*r*_*i*_ |*s*_*x*_) are the probability mass estimates of response probabilities across all trials and stimuli and conditioned on INS stimulus energy respectively. Probability mass functions (PMF) were estimated using histogram counts of spike responses with 1ms bin sizes for optimal information precision in A1 neurons(30). Calculations of MI from estimated PMFs was performed using the MIToolbox(31). Bias resulting from imperfect knowledge of true PMFs was corrected for using the method of quadratic extrapolation(32) as:

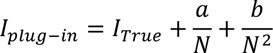

where *N* is the observed number of trials and *a*, *b* are free parameters dependent on stimulus-response relationships estimated by recalculating *I*_*plug−in*_(*R*; *S*_*x*_) at 50% and 25% of total samples and then performing least squares fit to the quadratic equation above. Information in spike trains was measured from onset of the stimulus till offset + 2 ms to account for offset responses.

#### 3.3 Joint Peri-stimulus Time Histogram Analysis

To assess spread of activation across cortical neurons, joint-peristimulus time histogram (JPSTH) analysis was performed(33). JPSTHs quantify dynamical, correlated activity between two neurons in response to a time-locked stimulus and thus represent purported functional connectivity from a source. JPSTHs were calculated using methods of Aertsen et al(34). Neurons were compared across each active electrode by first estimating joint densities of neuron PSTHs as:

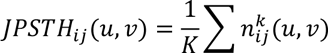

Where 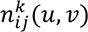 represents the spike count in bin *u*, *v* locked to stimulus repetition *k* for each neuron *i*, *j* for all stimulus repetitions *K*. The joint covariance due to co-stimulation of neurons from INS stimulation is calculated as the outer product of the PSTHs undertest

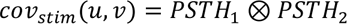

and represent stimulus-induced co-variation. Functional connectivity after stimulation was then calculated as

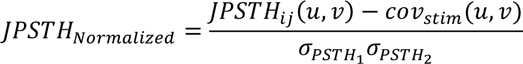

Distance of spread of activation was then calculated as the Euclidean distance between correlated neuron responses on each recording electrode given correlation between units. Animals receiving linear arrays (n=1) were excluded from this analysis as array geometry is not optimal for analyses assessing spread within cortical layers.

#### 3.4 Immunohistochemistry

At the end of experiments, rats were euthanized via a barbiturate overdose (beauthanasia 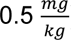) and underwent trans-cardiac perfusion of phosphate-buffered saline (PBS) and 0.4% paraformaldehyde. Brains were sliced into 20-50 *μm* slices using a cryotome and stored for immunohistochemistry. Brain slices containing the MGB were stained with NeuN (Abcam, Cambridge UK) conjugated to an Alexa-Fluor 488 secondary to label neurons and GFAP (Abcam) conjugated to an Alexa-Fluor 647 secondary to label reactive astrocytes. Full immunohistochemistry protocol is provided in the supplementary information. Slices were mounted and imaged using a Zeiss LSM710 confocal microscope (Zeiss, Jena GE) at 10x magnification resulting in an effective pixel size of 2.77*μm*^2. Tile scans across the length and height of the slice were made and stitched together using Zen10 (Zeiss) imaging suite. The MGB was identified via anatomical markers in conjunction with a rat stereotaxic atlas(35).

#### 3.5 Statistical Methods

Postsurgical changes in MLRs were assessed using the nonparametric Wilcoxon signed-rank test with comparisons made between pre and post implant first wave peak positivity (P1) representative of short latency brain stem responses, and first wave peak negativity (N1) and second peak positivity (P2) representative of later thalamocortical responses(36) (Fig 2A) with significance level set to *p* < 0.05.

To assess dose-response characteristics of thalamocortical recruitment from INS stimulation, a random effects multilinear regression model was utilized. Random effects repeated measures regression models were used to account for differences within subjects resulting from differences in recorded neuron physiology and distance from neuron to electrode as well as differences between subject responses. The multi-regression model was defined as

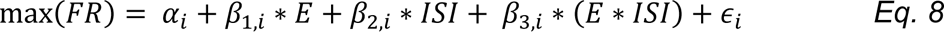

where FR is the BARS estimate of evoked firing rate, energy is the applied laser energy, ISI is the interstimulus interval, *i* is the mapping index codifying a neuron on an electrode of a given subject, and *ϵ* is estimated model error. The *α* coefficient corresponds to basal firing rate and β coefficients correspond to slope parameters of applied laser energy, ISI, and energy-ISI interactions. We chose to perform Bayesian inference to estimate model parameters because Bayesian methods are particularly powerful in modeling hierarchical random effects models(18, 37) and allow for robust and informative evaluation of regression parameters in posterior probability distributions. Parameter posterior distributions were summarized by their maximum a priori estimates (MAP), the most probable value and posterior 95% highest density intervals (HDI), quantifying the 95% most likely parameter values. Regression coefficients were considered significant if the 95% HDI did not overlap 0 in line with Bayesian inference convention(17). Bayesian formulations require choosing of prior distribution on regression parameters. As little is known about the effects of INS on thalamocortical firing patterns, normal distributions with a hyperprior variance of 5 were used. To ensure that prior distributions did not dominate observed data, prior distribution sensitivity analyses were performed (supplementary information (SI): Bayesian Model Specification). Analysis of observed data distributions suggested that log transformation of predictor laser parameters and evoked firing rate predictions provided best fit models. This was confirmed by post-hoc model comparisons and parameter sensitivity analyses (SI: Bayesian model specification).

Significance of stimulation and JPSTH correlations was assessed via shuffled permutation testing in which a null hypothesis was set such that *ρ*(*PSTH*_1_, *PSTH*_2_) = 0. For each pairwise correlation, one PSTH had bin counts shuffled following a uniform distribution and correlations recalculated. A total of 5000 shuffles for each pairwise comparison were performed. The number of shuffled correlations which were at least as extreme or greater than non-shuffled correlation was counted. Empirical p-values were calculated as the number of observed shuffles with correlation values greater than test correlations divided by number of permutation trials. Correlations were considered significant if empirical p-values < 0.05.

#### 3.6 Bayesian Multinominal Regression

Bayesian formulations of multinominal regressions were utilized to assess cortical neuron firing class dependence on stimulation parameters. Multinominal regression models were of the form

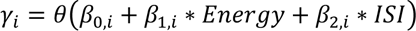

where *β*_0_ is the regression intercept term, *β*_1,2_ are the energy and ISI regression slope coefficients respectively, and *θ* is the softmax operator mapping regression to probability space corresponding to highest probability of class membership in class *γ*_*i*_. Due to indeterminacy of regression coefficients inherent to nominal models, predictions of class membership can only be made in reference to a reference category. To this end, we chose the reference category to be *onset+inhibition* which is the data category with the largest number of members in our dataset. Therefore, multinominal regressions are interpreted as the log odds of moving from the most populous class to a different firing class contingent upon INS parameters. Bayesian methods were used for parameter estimation and inference. Models were built in Python using PyMC v4. Prior distributions on *β* coefficients were chosen to be zero mean, variance 1 normal distributions which showed best fit to our observed data. Prior sensitivity analysis were performed and quantified in SI:Bayesian Model Description. Parameter posterior distributions were summarized by their maximum a priori estimates (MAP), the most probable value and posterior 95% highest density intervals (HDI), quantifying the 95% most likely parameter values. Regression coefficients were considered significant if the 95% HDI did not overlap 0 in line with Bayesian inference convention.

#### 3.7 PSTH classification via principle components analysis

Whether observed PSTHs formed distinct clusters of responses or exist across a continuum of the response space was assessed using principle components analysis. Individual PSTHs were represented as row vectors (*i*) = *PSTH*_*i*_ in a response matrix *r* = *N* × *b* where N is the total number of evoked PSTHs and b is the total number of bins. PSTHs were constructed from 5 ms bins. Response matrix r was then decomposed into a vector of principle components *r* with weights *w ϵ*{0,1}. Finding components of maximal variance was found by ensuring the first weight component (1) satisfies

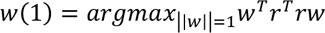

The remaining *k* − 1 components and weights are then estimated as

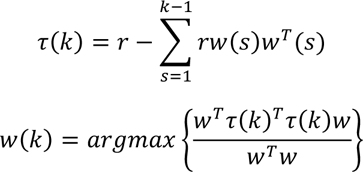

The first 3 components representing 65.11% of explained variance were extracted for clustering. Components were then mapped to response classes for visualization.

### 4 Immunohistochemistry Protocols

#### 4.1

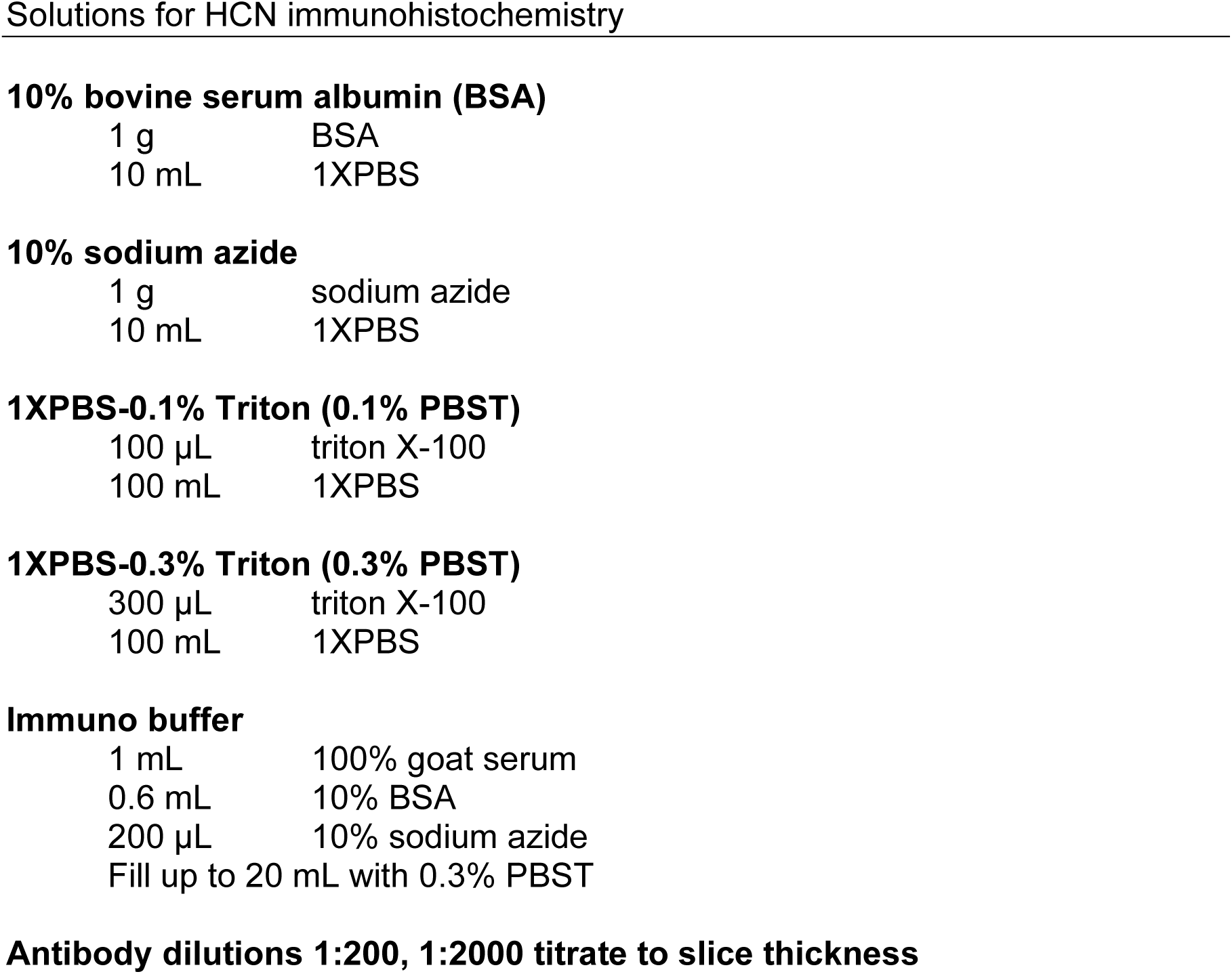

**10% bovine serum albumin (BSA)**
|  |  |
| --- | --- |
| 1 g | BSA |
| 10 mL | 1XPBS |
| **10% sodium azide** |  |
| 1 g | sodium azide |
| 10 mL | 1XPBS |
| **1XPBS-0.1% Triton (0.1% PBST)** |  |
| 100 µL | triton X-100 |
| 100 mL | 1XPBS |
| **1XPBS-0.3% Triton (0.3% PBST)** |  |
| 300 µL | triton X-100 |
| 100 mL | 1XPBS |
| **Immuno buffer** |  |
| 1 mL | 100% goat serum |
| 0.6 mL | 10% BSA |
| 200 µL | 10% sodium azide |
| Fill up to 20 mL with 0.3% PBST |  |

**Antibody dilutions 1:200, 1:2000 titrate to slice thickness**

#### 4.2 IHC Protocol

| Serum |  |  |  |  |
| --- | --- | --- | --- | --- |
| Primary | Host | Concentration | Secondary | Concentration |

Protocol:
1. 2x 5-min wash in 1XPBS-0.1% Triton (0.1% PBST) at room temperature. 2. 30-min incubation in 1XPBS-3% Triton (0.3% PBST) at room temperature. 3. 1-hour incubation in immuno buffer. 4. 48-hour incubation in 1° antibody diluted with immuno buffer at 4°C. 5. 3x 10-min washes in 0.1% PBST.

## 5 Supplementary Figures

**Figure S1:**
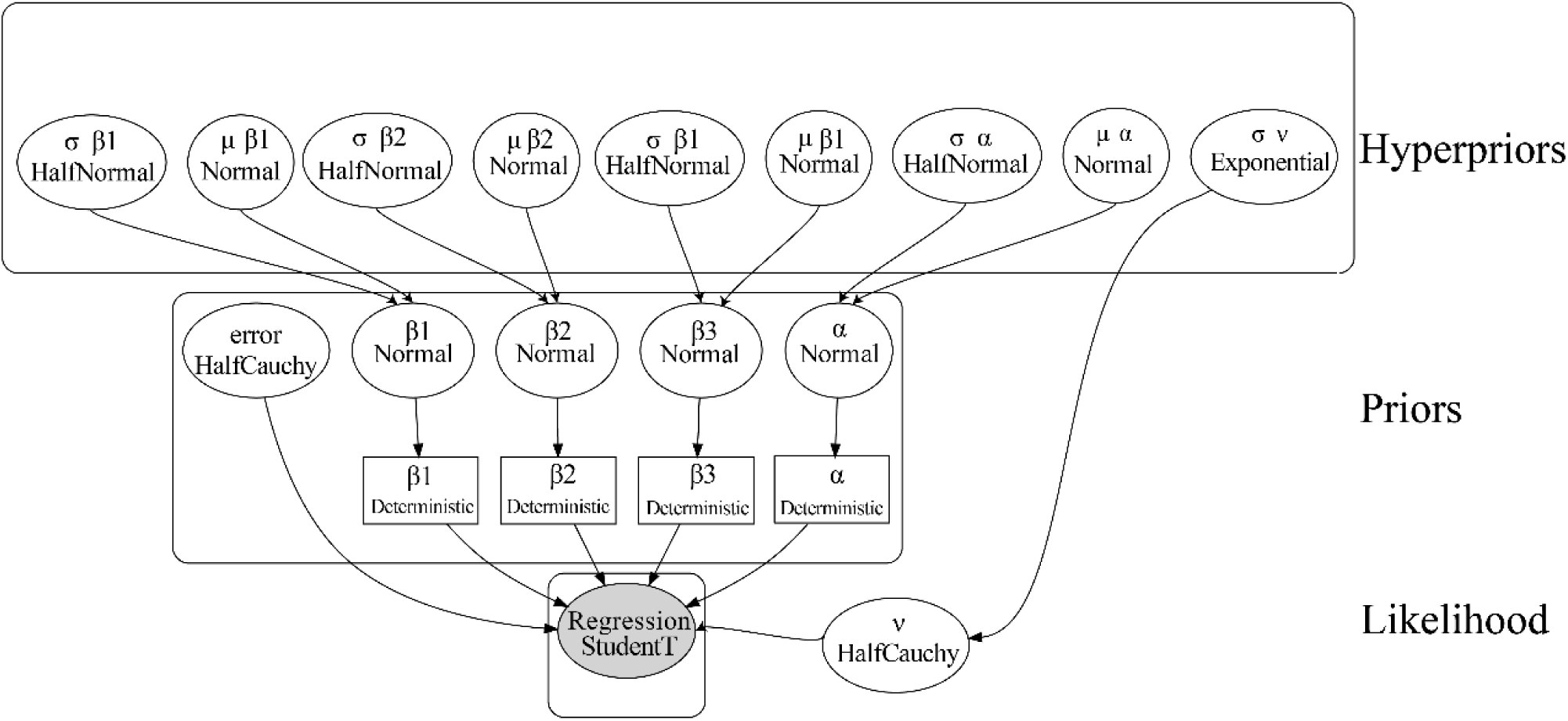
Schematic of Bayesian hierarchical Multilinear regression utilized in this study. Deterministic nodes were included in the model to prevent MCMC sampling from entering regions of solution space which are difficult to move away from.

**Figure S2:**
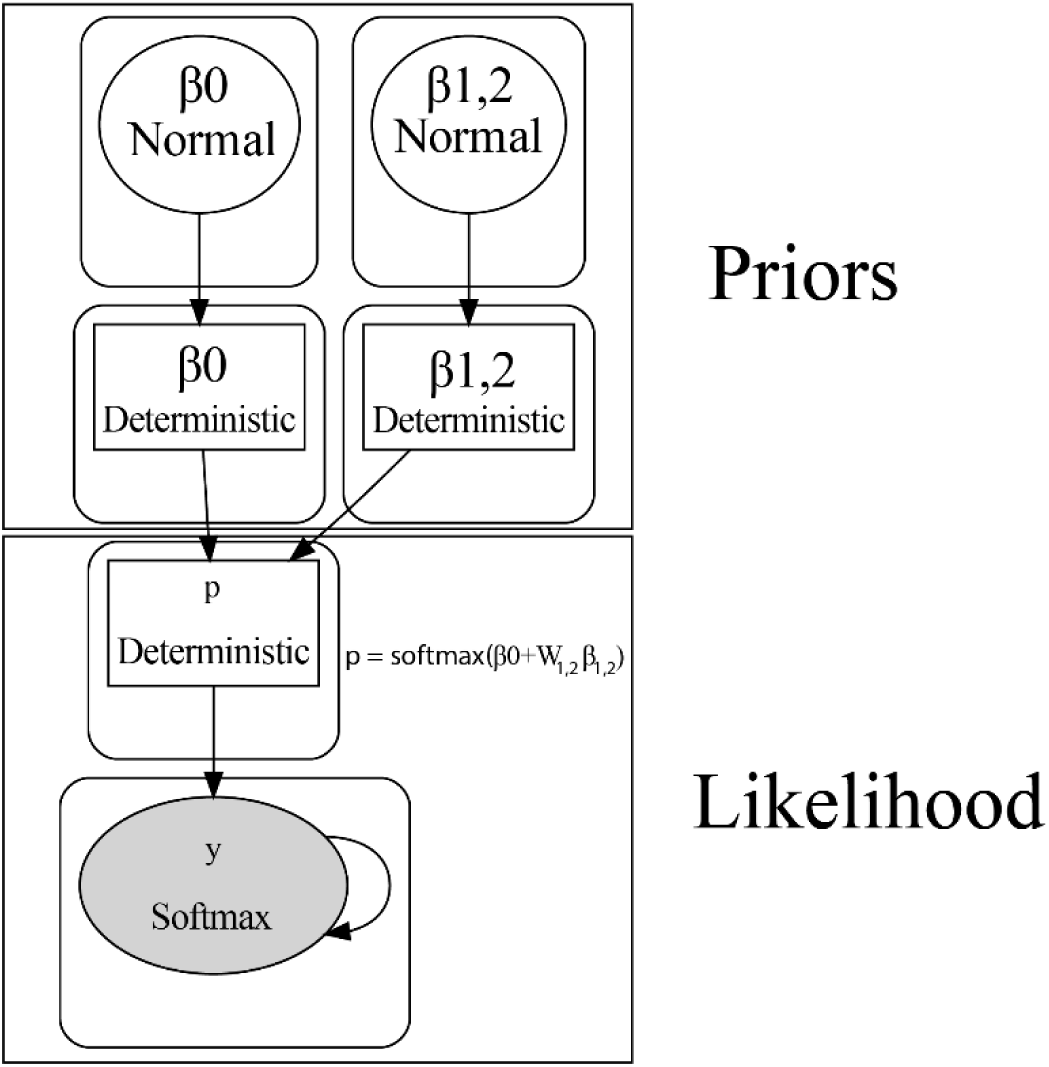
Schematic of Bayesian multinominal regression utilized in this study. Deterministic nodes were included in the model to prevent MCMC sampling from entering regions of solution space which are difficult to move away from.

**Figure S3:**
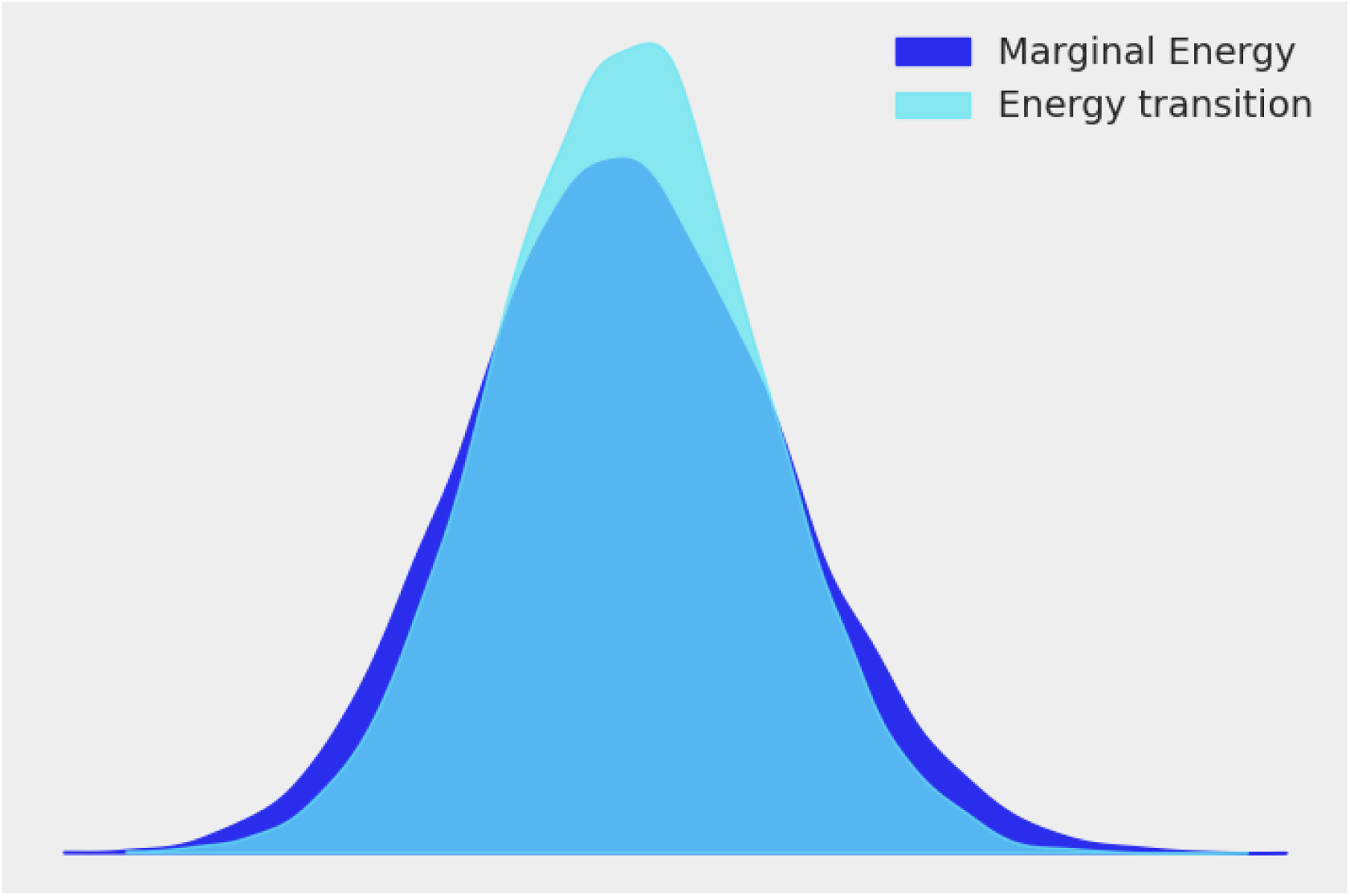
Energy trace of NUTS trace for the Bayesian hierarchical regression model.

**Figure S4:**
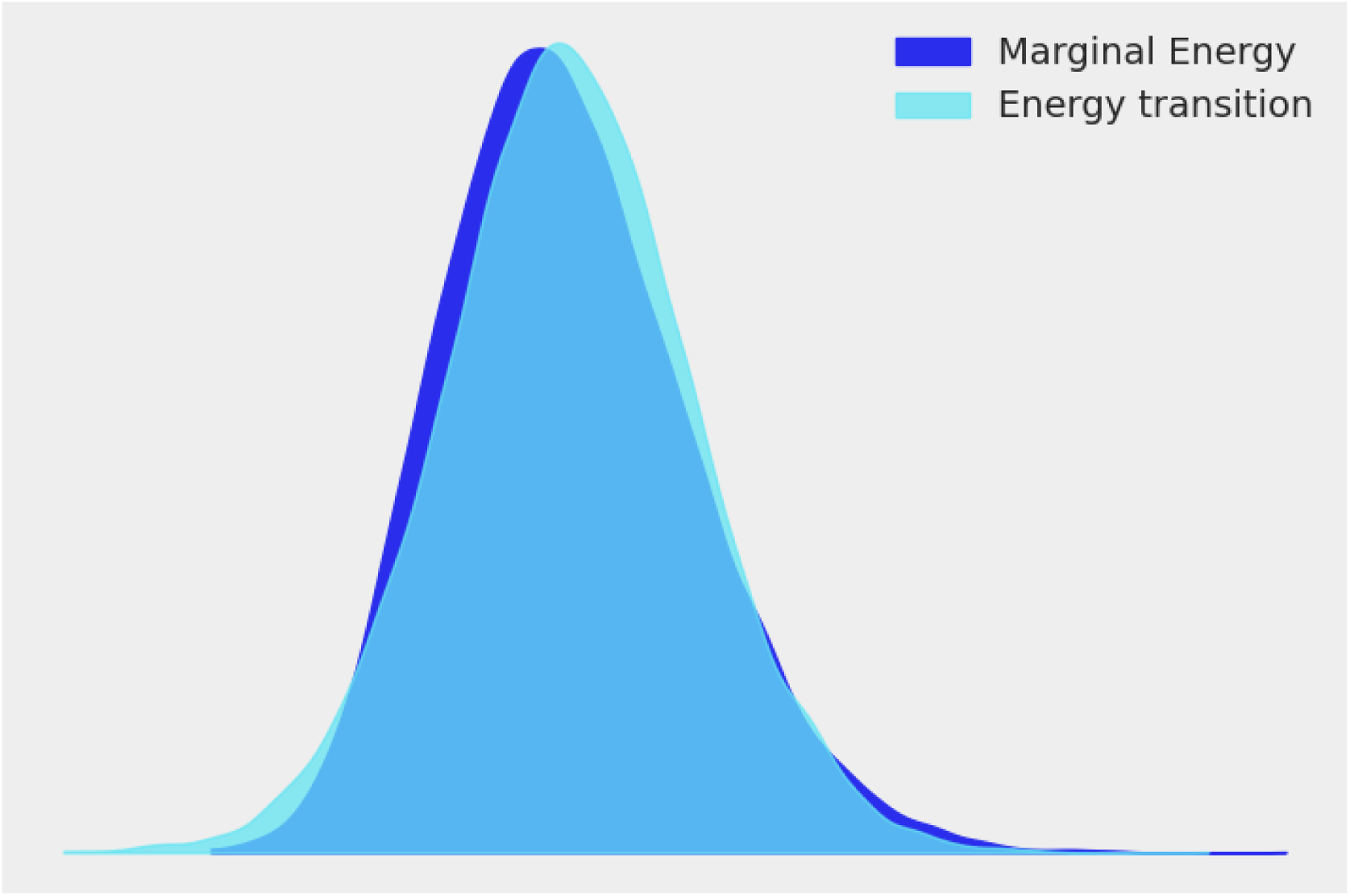
Energy trace of NUTS trace for the Bayesian multinominal regression model.

**Figure S5:**
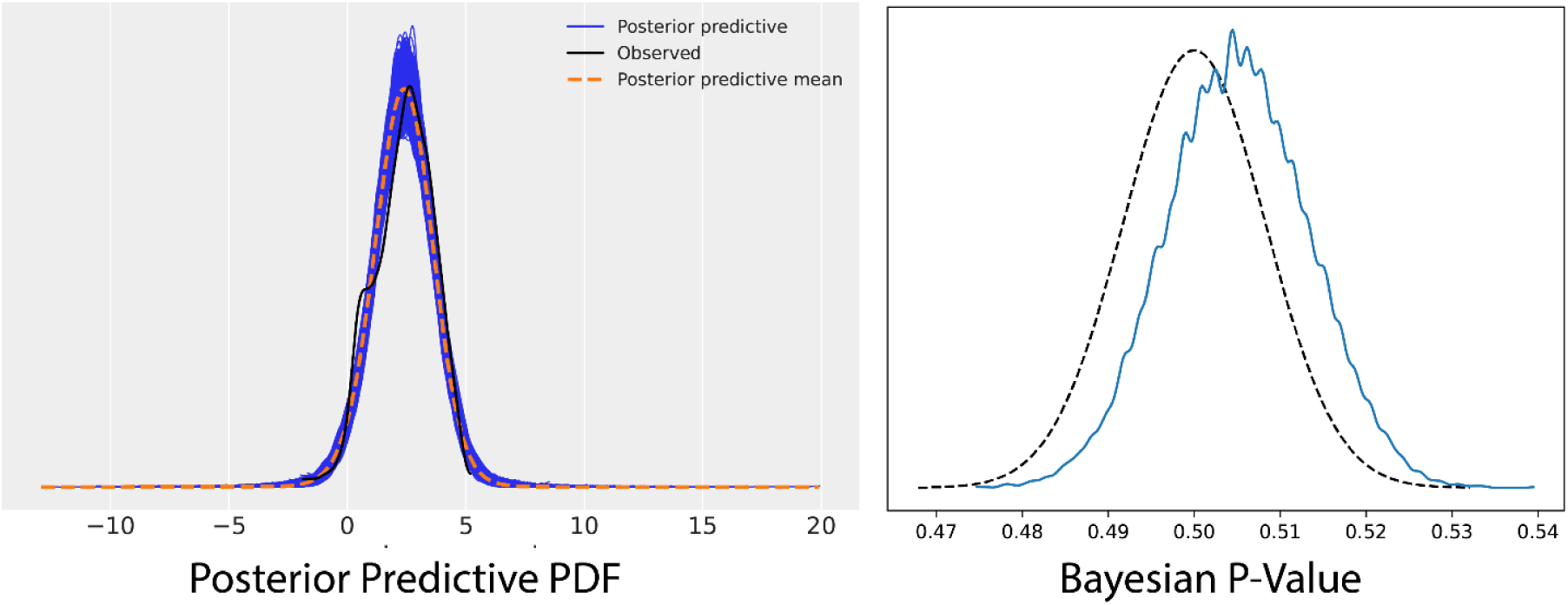
Posterior predictive checks for the Bayesian hierarchical multilinear regression models. Strong overlap of posterior predictive distribution with observed density estimates and p-values near 0.50 indicate model was well fit to observed data.

**Figure S6:**
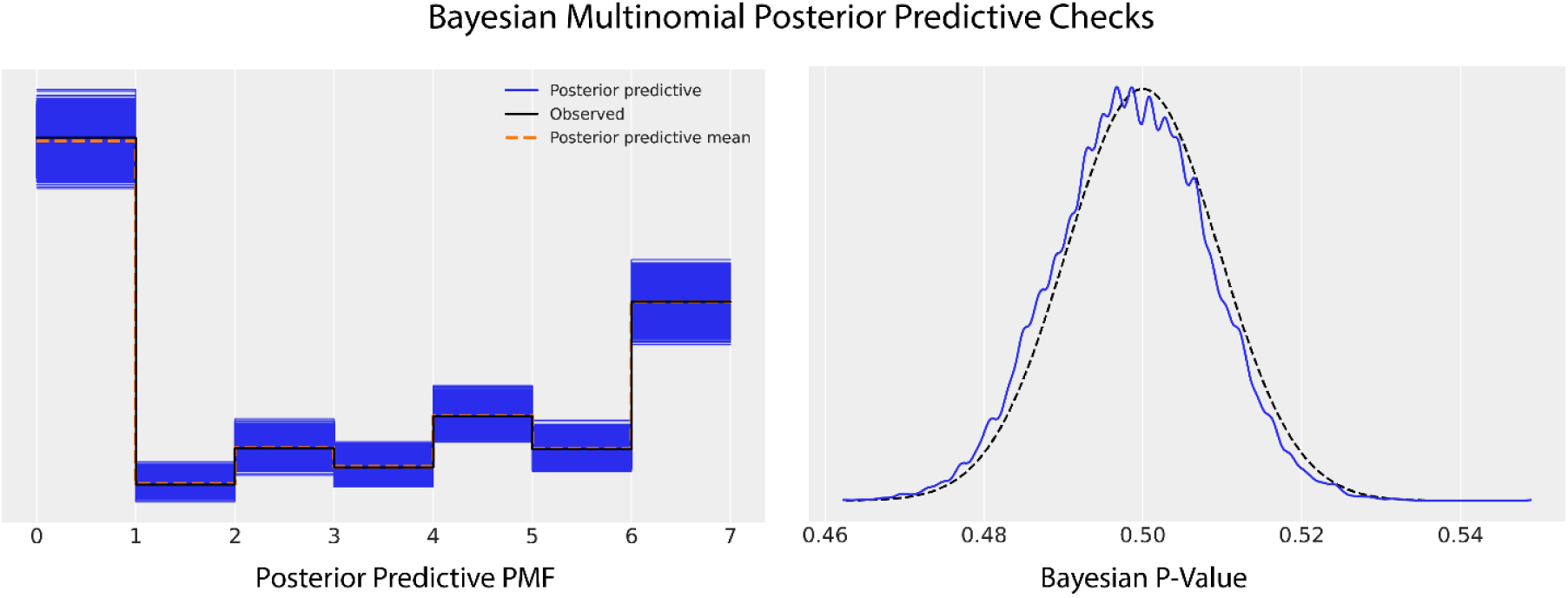
Posterior predictive checks for the Bayesian multinominal regression models. Strong overlap of posterior predictive distribution with observed probability mass estimates and p-values near 0.50 indicate model was well fit to observed data.

**Figure S7:**
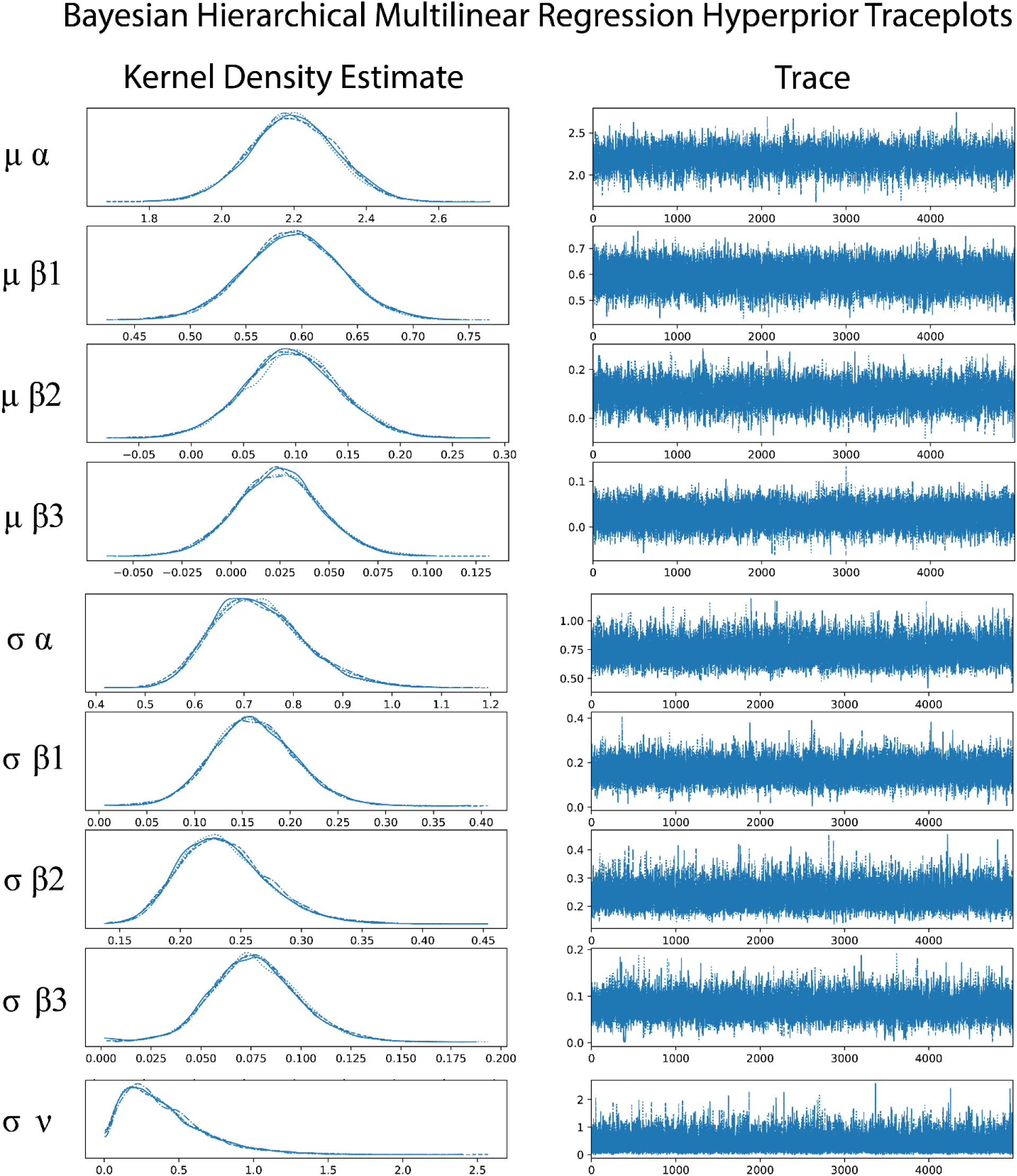
Traceplots and kernel density estimates for hyperpriors of the Bayesian hierarchical multilinear regression model.

**Figure S8:**
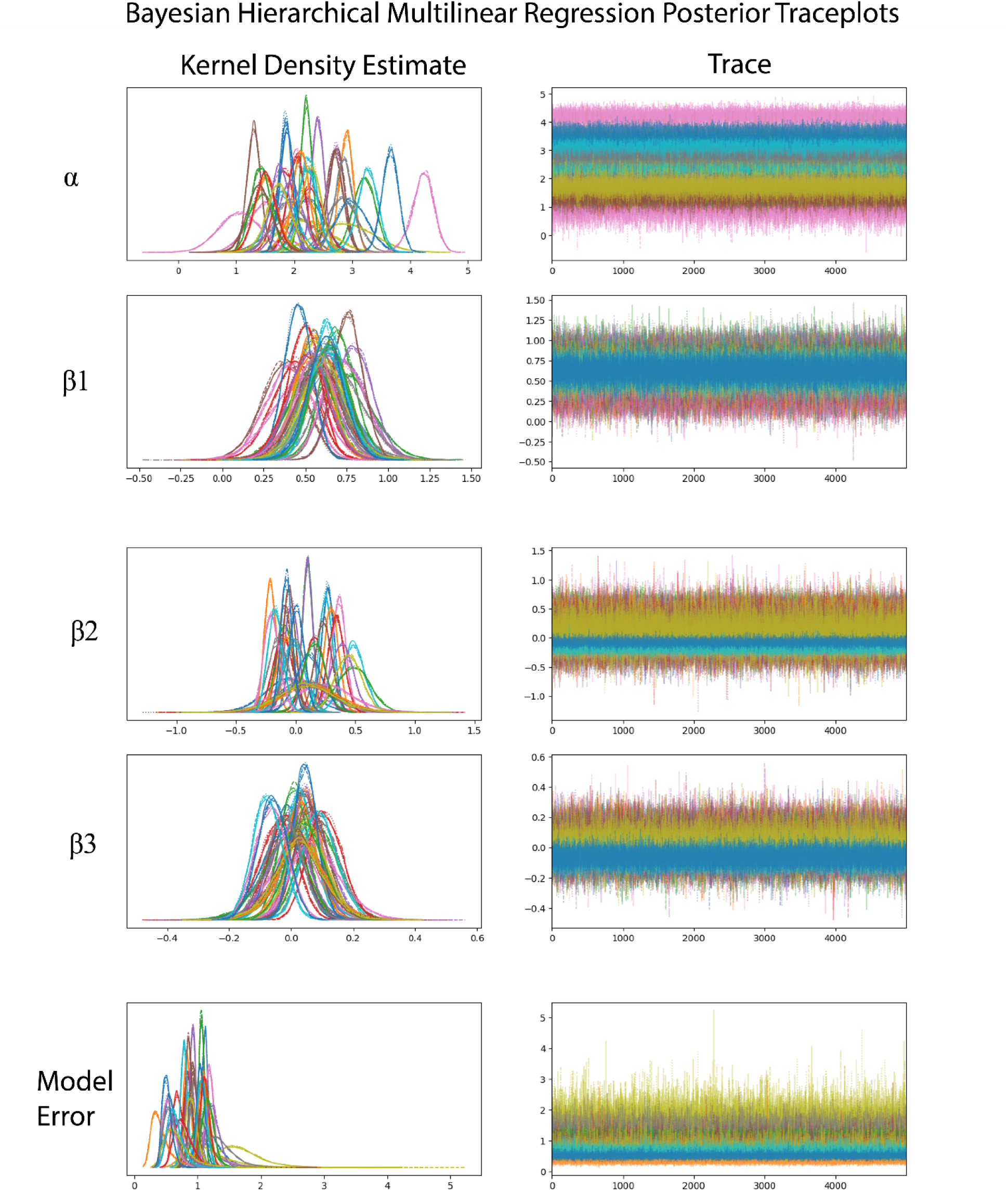
Traceplots and kernel density estimates for posteriors of the Bayesian hierarchical multilinear regression model. Model is partially pooled, with each color representing estimation of the parameter for a given neuron on a given site.

**Figure S9:**
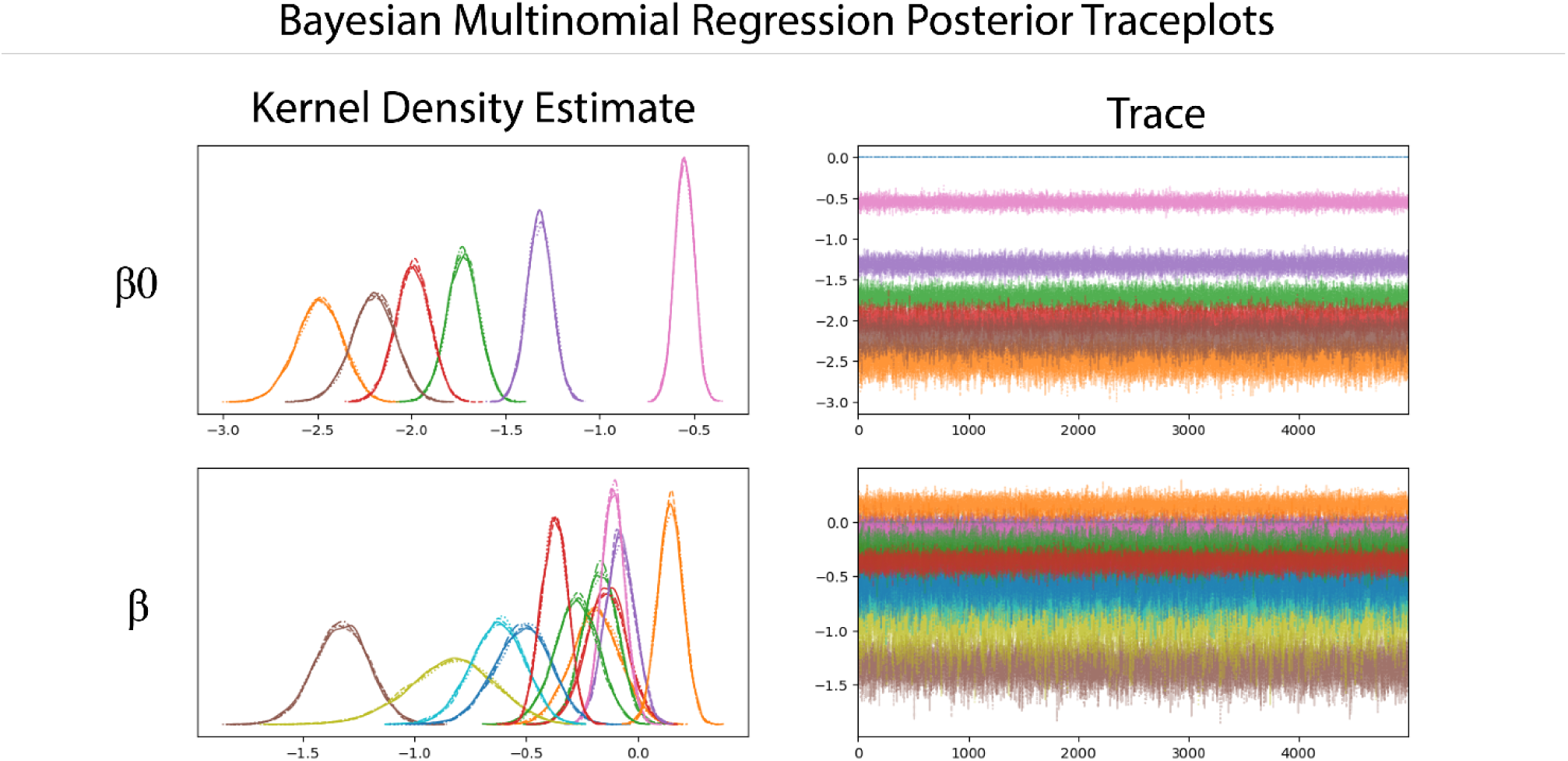
Traceplots and kernel density estimates for posteriors of the Bayesian multinominal regression model. Each color represents estimation of the parameter for a given firing class. As *β* was cast to encapsulate regression coefficients for both laser energy and ISI (see model description), both coefficients are presented on the same kernel density and trace plot.

**Figure S10:**
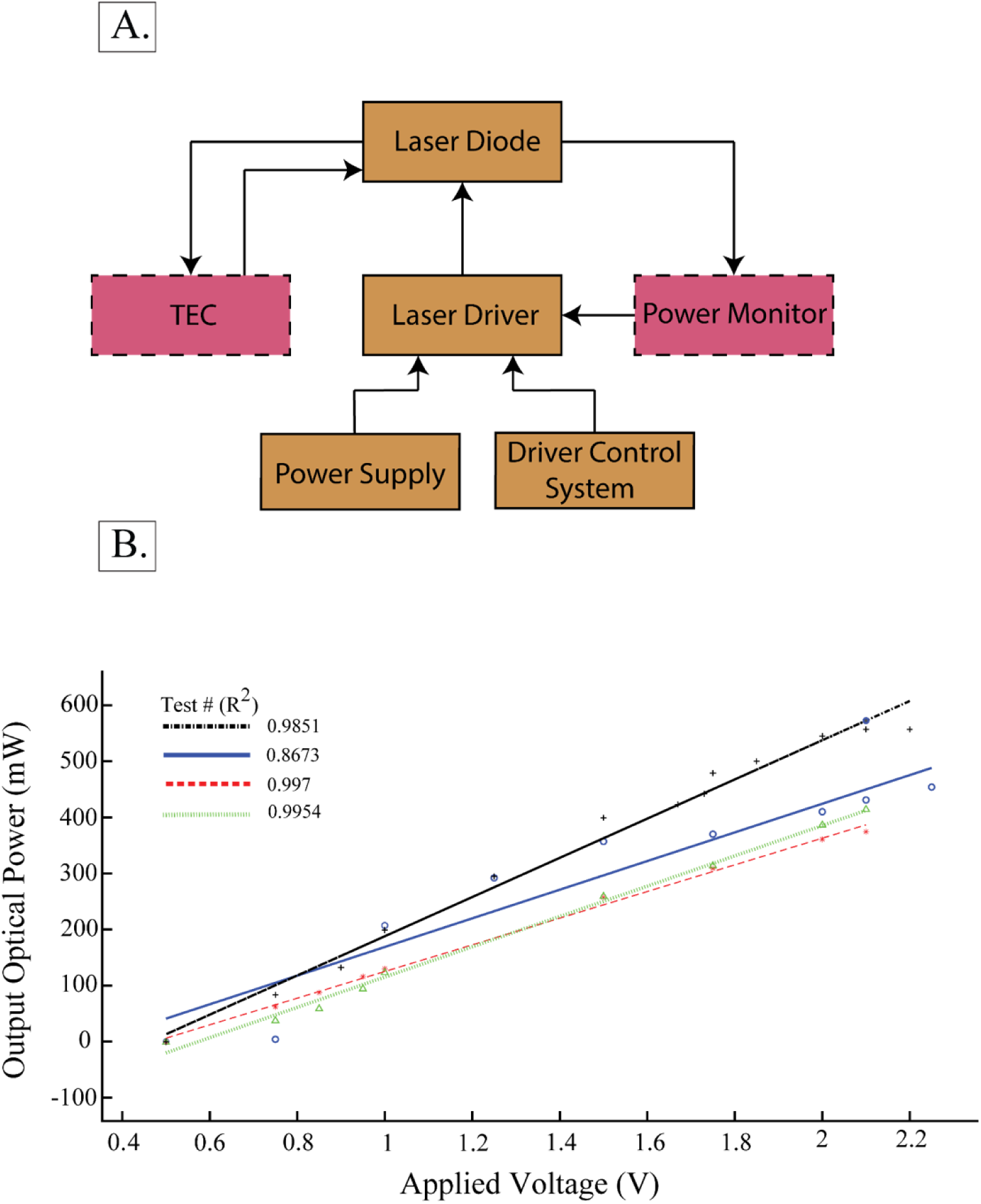
INS laser system description and validation. A.) Block diagram detailing system composition. Boxes colored gold consist of required hardware necessary for laser operation. Boxes colored pink with dotted lines are optional, but strongly suggested control modules to accompany primary system design. B.) Measured output optical power in response to open loop application of voltage pulses to the laser driver shows strong output linearity with respect to applied modulation voltage. Four calibration curves taken over the span of a year show the need for routine calibration and/or the use of a power control system in closed loop operation. N.B. Applied voltage refers to the voltage applied from laser control hardware, not the voltage at the laser diode. In our case, control voltages were generated from a TDT RZ-2 analog output.

**Figure S11:**
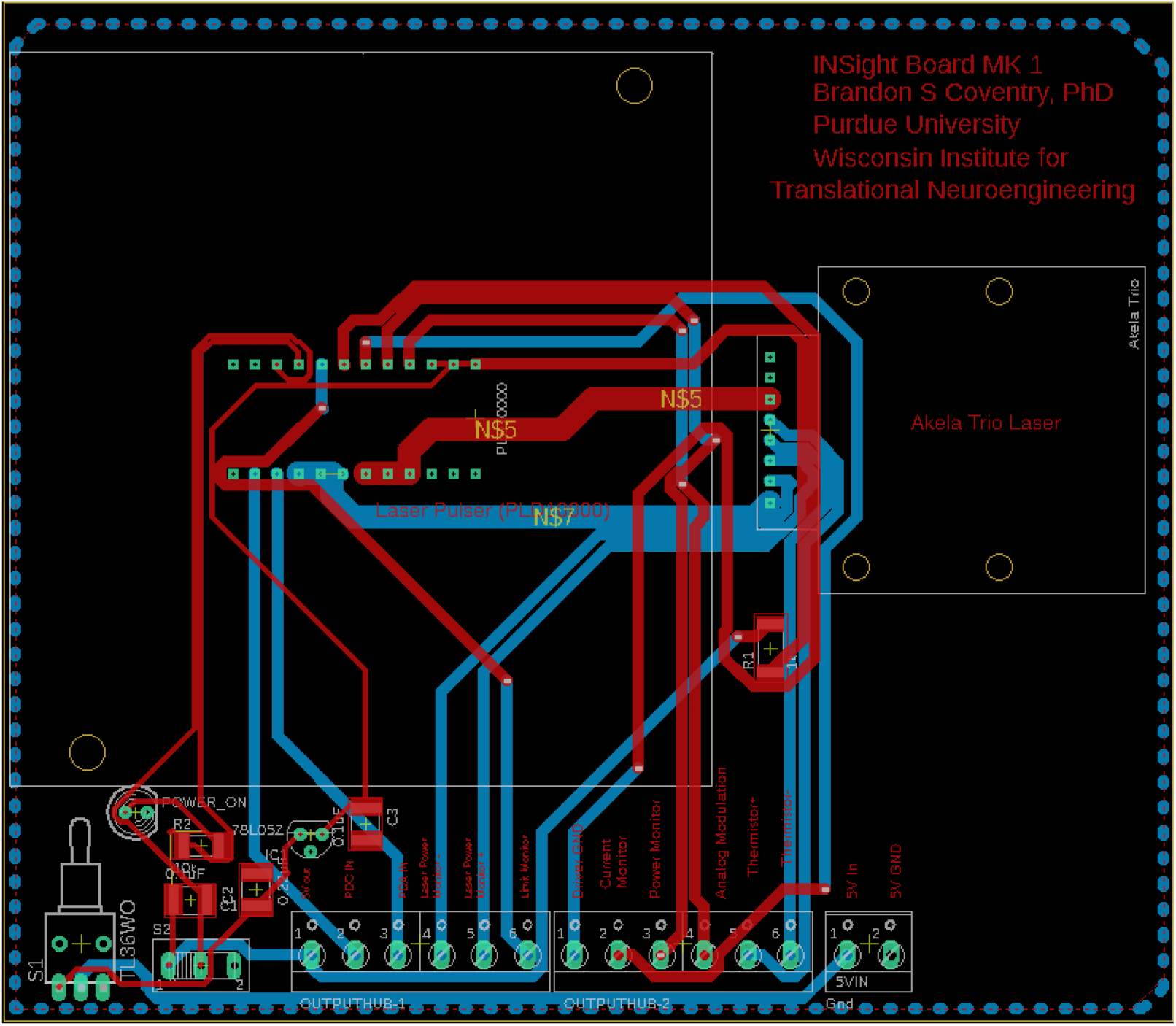
Example laser driver and diode printed circuit board interface. Ground layers have been removed for image clarity. Pads associated with the Akela Trio laser diode should be populated with pogo pins and secured by screwing in the diode through the predrilled screw holes. All Gerber files are available at https://github.com/bscoventry/INSight.

**Figure S12:**
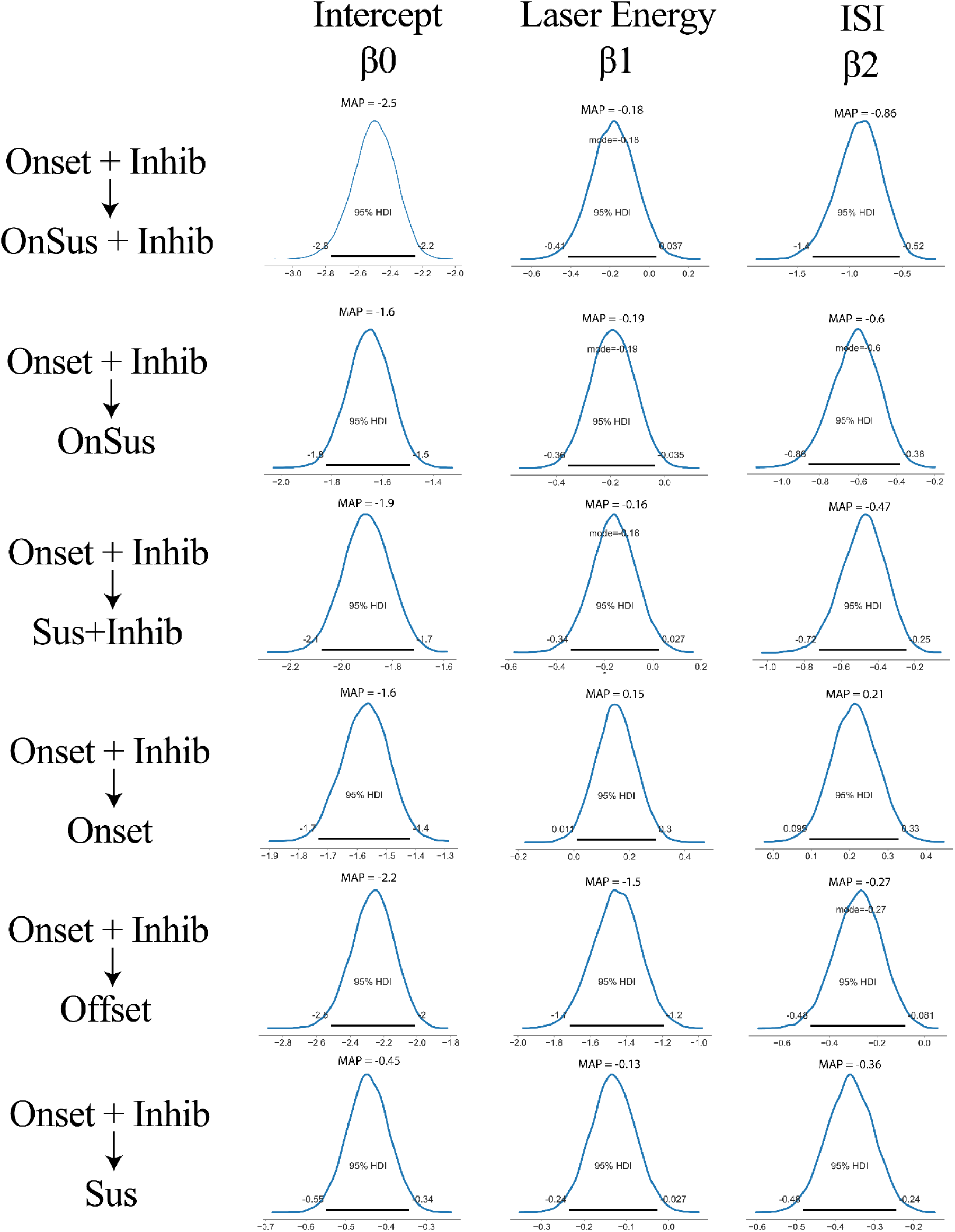
Posterior distributions of multinominal regression parameters suggest that evoked cortical firing classes are partly influenced by stimulation parameters. The most populous class onset+inhib was chosen for the reference class. Class membership changes resulting from changes in laser parameters was considered significant if 95% HDI did not overlap 0. Movement from membership of onset+inhib to onset arose from increasing laser energy and ISI. Movement from onset+inhib to onset-sustained resulted from decreased laser energy and decreased ISIs while movement of onset-inhib to onset-sustained+inhib was due only to decreases in ISI. Likewise, movement from onset+inhib to sustained firing resulted from slight decreases in applied laser energy but larger decreases in ISI while movement towards sustained+inhibition was marked by larger decreases in ISI. Finally movement to offset responses was marked by larger decreases in laser energy and smaller decreases in laser pulse ISI. This data taken together suggests a complex interplay between stimulation parameters and native cellular biophysics.

**Figure S13:**
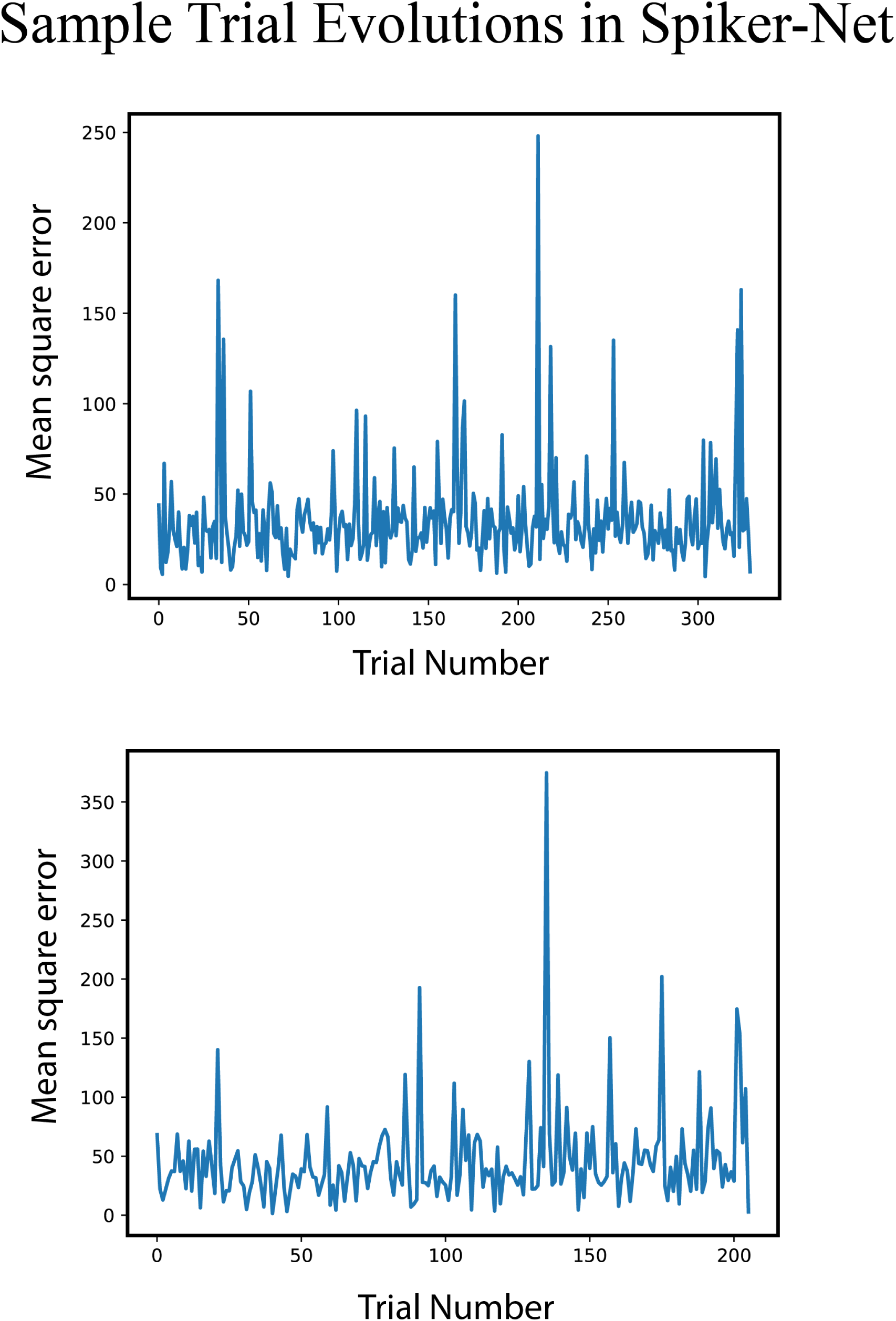
Plotting SpikerNet mean-square error during training reveals searching and targeting behavior, with periods marked by large mean-square errors indicative of algorithmic searching behavior followed by targeting optimal stimuli as evidenced by low mean-square error.

## 6 Software and Data Repositories

### 6.1 Dataset S1

Datasets can be found in the following open science framework repository: https://osf.io/w4ufh/

### 6.2 Software S1

Analysis programs can be found at the following github repository: https://github.com/bscoventry/OpticalTCNeuromodulation

INSight design files and software can be found at the following github repository: https://github.com/bscoventry/INSight

